# Cell-type-specific meQTL extends melanoma GWAS annotation beyond eQTL and informs melanocyte gene regulatory mechanisms

**DOI:** 10.1101/2021.03.23.436704

**Authors:** Tongwu Zhang, Jiyeon Choi, Ramile Dilshat, Berglind Ósk Einarsdóttir, Michael A Kovacs, Mai Xu, Michael Malasky, Salma Chowdhury, Kristine Jones, D Timothy Bishop, Alisa M Goldstein, Mark M Iles, Maria Teresa Landi, Matthew H Law, Jianxin Shi, Eiríkur Steingrímsson, Kevin M Brown

## Abstract

While expression quantitative trait loci (eQTL) have been powerful in identifying susceptibility genes from genome-wide association studies (GWAS) findings, most trait-associated loci are not explained by eQTL alone. Alternative QTLs including DNA methylation QTL (meQTL) are emerging, but cell-type-specific meQTL using cells of disease origin has been lacking. Here we established an meQTL dataset using primary melanocytes from 106 individuals and identified 1,497,502 significant *cis*-meQTLs. Multi-QTL colocalization using meQTL, eQTL, and mRNA splice-junction QTL from the same individuals together with imputed methylome-wide and transcriptome-wide association studies identified susceptibility genes at 63% of melanoma GWAS loci. Among three molecular QTLs, meQTLs were the single largest contributor. To compare melanocyte meQTLs with those from malignant melanomas, we performed meQTL analysis on skin cutaneous melanomas from The Cancer Genome Atlas (n = 444). A substantial proportion of meQTL probes (45.9%) in primary melanocytes are preserved in melanomas, while a smaller fraction of eQTL genes is preserved (12.7%). Integration of melanocyte multi-QTL and melanoma meQTL identified candidate susceptibility genes at 72% of melanoma GWAS loci. Beyond GWAS annotation, meQTL-eQTL colocalization in melanocytes suggested that 841 unique genes potentially share a causal variant with a nearby methylation probe in melanocytes. Finally, melanocyte *trans*-meQTL identified a hotspot for rs12203592, a *cis*-eQTL of a transcription factor, IRF4, with 131 candidate target CpGs. Motif enrichment and IRF4 ChIPseq analysis demonstrated that these target CpGs are enriched in IRF4 binding sites, suggesting an IRF4-mediated regulatory network. Our study highlights the utility of cell-type-specific meQTL.

## Introduction

Expression quantitative trait loci (eQTL) studies have been powerful for nominating candidate causal genes for loci identified via genome-wide association studies (GWAS) of many complex traits and diseases, including cancer susceptibility. Most prominently, the Genotype-Tissue Expression (GTEx) project has made eQTL data publicly available for more than 50 tissue types^1^. Most eQTL datasets including GTEx, however, are based on heterogeneous bulk tissues, where cell-type-specific allelic regulation of gene expression in rarer cell types may be obscured by signals from other cell types, and thus may go undetected. Colocalization analyses using the most recent GTEx dataset demonstrated that a median of 21% of GWAS loci from 87 tested complex traits colocalized with a *cis*-eQTL when aggregated across 49 tissue types^1^. While cell-type interacting eQTLs by computational deconvolution of bulk tissue data improves colocalization compared to that by standard eQTL only^2, 3^, most GWAS loci nonetheless lack colocalizing eQTLs.

A recent melanoma GWAS meta-analysis identified a total of 54 loci reaching genome-wide significance^4^, increasing the total number of melanoma risk-associated loci by more than three-fold compared to the largest existing study^5^. We previously demonstrated that eQTLs from cultured melanocytes^6^, the cell type of origin for melanoma, efficiently identified candidate susceptibility genes for 25%^5, 6^ and 16%^4^ of two recent melanoma GWAS loci through colocalization. Notably, as melanocytes represent only a small fraction of typical skin biopsies, even this moderately sized melanocyte eQTL dataset (n = 106) was able to identify candidate causal genes that were not captured by GTEx skin tissue eQTLs from sample sets three times larger^6^. These data highlighted the utility of cell-type-specific QTL resources, however, eQTL alone was still not sufficient to explain the majority of GWAS loci.

DNA methylation of cytosine at CpG dinucleotides is an important mode of epigenetic gene regulation. While CpG methylation is interconnected with mRNA expression, their relationship is rather complex. In tumors, hypermethylation has been observed in the promoters of inactivated tumor suppressor genes^7^. Gene body methylation, on the other hand, is usually correlated with higher mRNA expression and tends to be inversely correlated with promoter methylation^8^. Further, it is not always clear whether methylation/demethylation actively initiates gene expression repression/activation, or instead, methylation levels reflect repressed/activated expression status^9^. While DNA methylation has been more widely studied as a marker of epigenetic regulation in population studies (e.g. EWAS^10^), DNA methylation is also under tight genetic control as shown by methylation QTL (meQTL) studies. In particular, *trans*-meQTL has been powerful in identifying transcription factor-mediated regulation networks and large numbers of target CpGs^11–13^, in contrast to relatively small numbers of *trans*-eQTL genes or *trans*-sQTL genes when using gene expression data^1^.

meQTL studies to date have largely been limited to blood and blood-related cell types^11, 12, 14–20^, with a few exceptions of studies of normal bulk tissues^13, 21–23^ and tumor tissues^24, 25^. Overall, cell-type-specific meQTL studies from non-blood samples have largely been lacking. Particularly in the context of cancer, understanding a heritable component of DNA methylation in the cell types where the tumor originates may help answer questions about how methylation and gene expression are co-regulated through genetic variants and how much of that genetic regulation is still observed during the malignant transformation where multiple genetic and non-genetic events could mask gene expression variance explained by germline variants.

In this study, we explore the roles of cell-type-specific meQTLs derived from human primary melanocytes in explaining melanoma-risk associated genetic signals through multi-QTL colocalization as well as imputed methylome-wide association study (MWAS)^26^. We further compare genetic control of DNA methylation in melanocytes with that of malignant melanoma tissues. We then interrogate the relationship of eQTL and meQTL in melanocytes and further identify a melanocyte-specific transcriptional hub through *trans*-meQTL study.

## Material and Methods

### DNA methylation profiling

Genome-wide DNA methylation was profiled on Illumina HumanMethylation450 BeadChip (Illumina, San Diego, USA). Genomic DNA was extracted from primary cultures of melanocytes from 106 newborn males mainly of European descent as previously described^6^, and DNA methylation was measured according to Illumina’s standard procedure at Cancer Genomics Research Laboratory (CGR), National Cancer Institute. Basic intensity QC was performed using the minfi R package^27^. Briefly, raw methylated and unmethylated intensities were background-corrected, and dye-bias-equalized to correct for technical variation in signal between arrays. The following criteria were applied to filter probes and samples: 1) Probes located on chrX and chrY were removed. 2) Probes including common SNPs with minor allele frequency (MAF) > 5% (1000 Genomes, phase 3, EUR) were removed. No melanoma GWAS loci were found within 1 Mbp of these SNPs. 3) Probes located in repetitive genomic regions (repeatmask hg19 database) were removed. 4) Probes with detection *P*-value >0.01 were marked as missing. Probes with a missing rate >5% were removed and samples with a missing rate >4% were removed. 5) Control samples and samples without matched genotyping data were removed. 6) For duplicated samples, the better one of the two was selected based on probe intensity, SNP call rate, and the percentage of missing probes. No batch effects or plating issues were identified across plates, wells, and barcode IDs based on the assessment of methylated and unmethylated intensities, failed samples, and beta distributions. Functional normalization implemented in the minfi R package^27^ was used to calculate the final methylation levels (beta value) after normalization. In total, we retained 386,520 probes and 106 samples for the downstream meQTL analysis. In addition, we calculated the top 10 probabilistic estimation of expression residuals (PEER)^28^ as the potential hidden covariates for QTL analysis.

### Quantification of RNA splicing

RNA-Seq data of the same 106 melanocytes from our previous publication^6^ were re-analyzed to quantify RNA splicing. The processed BAM files were used to create the junction files and intron clustering based on the instruction of LeafCutter^29^. The normalized quantification of 117,570 junctions was generated as the phenotype and 10 Principal Components (PCs) were included as covariates for splice QTL (sQTL) analysis.

### meQTL and sQTL detection

*Cis*-meQTL and *cis*-sQTL analyses were performed using the same *cis*-QTL pipeline and the same processed genotype data (vcf format) as described in our previous *cis*-eQTL analysis^6^. Briefly, FastQTL was used to perform *cis*-QTL mapping^30^, and nominal *P*-values were generated for genetic variants located within ±1 Mb of the transcription start sites (TSSs) for each probe or junction tested. For covariates of QTL analyses, we included 3 PCs inferred based on genotype data, and independent methylation variables (Pearson correlation coefficient < 0.8) from 10 PEER factors (meQTL), or independent splice junction usage variables (Pearson correlation < 0.8) from 10 PCs (sQTLs). The beta distribution-adjusted empirical *P*-values from FastQTL were then used to calculate *q*-values^31^, and a false discovery rate (FDR) threshold of ≤ 0.05 was applied to identify probes or junctions with a significant QTL (“meProbes” or “sJunctions”). We used a similar method as that for GTEx study (using FastQTL) to identify all significant variant-probe or junction pairs. In summary, a genome-wide empirical *P*-value threshold, *p_t_*, was defined as the empirical *P*-value of the probe or junction closest to the 0.05 *FDR* threshold. *P_t_* was then used to calculate a nominal *P*-value threshold for each gene based on the beta distribution model of the minimum *P*-value distribution *f(pmin)* obtained from the permutations for the probe or junction. Specifically, the nominal threshold was calculated as *F^-1^(p_t_)*, where *F^-1^* is the inverse cumulative distribution. For each probe or junction, variants with a nominal *P-value* below the probe or junction-level threshold were considered significant and included in the final list of genome-wide significant *cis*-QTL variants. The effect (slope) of QTLs is relative to the alternative allele.

### *trans*-meQTLs detection

Identification of *trans*-meQTLs has been described previously by Shi and colleagues^13^. Prior to meQTL analysis, each methylation trait was regressed against batches and independent PEER factors based on methylation profiles. The regression residuals were then quantile-normalized to the standard normal distribution *N*(0,1) as traits. The genetic association testing was performed using tensorQTL^32^, adjusted for the top three PCA scores based on GWAS SNPs to control for potential population stratification. To identify the threshold for genome-wide significant *trans*-meQTLs, the following statistical steps were applied. For each CpG probe, the *trans* region was defined as being more than 5 Mb from the target CpG site in the same chromosome or on different chromosomes. For the *n*th methylation trait with *m* SNPs in the *trans* region, let (*qn1*,*⋯*,*qnm*) be the *P*-values for testing the marginal association between the trait and the *m* SNPs. Let *pn* = min(*qn1*,*⋯*,*qnm*) be the minimum *P*-value for *m* SNPs and converted *pn* into genome-wide *P*-value *Pn* by performing one million permutations for SNPs in the *trans* region. As a *cis* region is very short compared with the whole genome, *Pn* computed based on SNPs in *trans* regions is very close to that based on permutations using genome-wide SNPs. Thus, we use the genome-wide *P*-value computed based on all SNPs to approximate *Pn*. Furthermore, all quantile-normalized traits follow the same standard normal distribution *N*(0,1); thus the permutation-based null distributions are the same for all traits. We then applied the Benjamini–Hochberg^33^ procedure to (*P1*,⋯,*PN*) to identify *trans*-meQTLs by controlling FDR at 1%, which corresponded to a nominal *P*-value of 1.03E-11.

### TCGA SKCM meQTL analysis

Four hundred and forty-four Skin Cutaneous Melanoma (SKCM) samples from The Cancer Genome Atlas (TCGA) with both genotype data and methylation data were included in our study. For genotype data, we collected our previously processed genotype data in vcf format^6^. The original raw intensity idat files from Human Methylation 450 array with matched genotype data were downloaded from NCI Genomic Data Commons Data Portal (GDC Legacy Archive, https://portal.gdc.cancer.gov). The same DNA methylation processing pipelines for melanocytes described above were applied to TCGA methylation data, which included 384,273 high-quality probes for the downstream analysis. We selected the 3 PCs calculated from genotype data and uncorrected 10 Peer Factors from methylation data for the meQTL analysis. In addition, we adjusted the copy number alterations for each probe by including the segmentation’s logR value as a covariate for meQTL analysis. The segmentation CNV data was calculated from the SNP array as TCGA level 3 dataset, which was collected from GDC portal. We followed the same melanocyte *cis*-meQTL analysis pipeline for the TCGA SKCM meQTL analysis. For *trans*-meQTL in TCGA SKCM, we only tested the association of significant melanocyte *trans*-meQTLs and applied a similar genome-wide *P*-value threshold (1.03E-11) between SNPs and distant CpG Probes.

### Pairwise meQTL sharing between primary melanocytes and TCGA SKCM

To test the sharing of all significant SNP-CpG probe pairs of our melanocyte *cis*-meQTLs with those identified in TCGA SKCM, we calculated pairwise π1 statistics, where π1 is the proportion of all genome-wide significant meQTLs (using a threshold of *FDR* < 0.05) from one dataset found to also be genome-wide significant in the other. We used QVALUE^31^ to calculate π1, which indicates the proportion of true positives. A higher π1 value indicates an increased replication of meQTLs.

### Multi-QTL colocalization

Melanoma GWAS summary statistics from a meta-analysis of 36,760 clinically confirmed and self-reported cutaneous melanoma cases were collected from a recent study^4^, which included 54 significant loci with 68 independent SNPs. All study participants provided informed consent reviewed by IRBs, including 23andMe participants with online informed consent and participation, under a protocol approved by the external AAHRPP-accredited IRB, Ethical & Independent Review Services (E&I Review). We performed multi-QTL colocalization analyses among GWAS, eQTL, meQTL, and sQTL datasets. HyPrColoc^34^ was used to perform colocalization analysis with the default parameters: prior.1 (1e-4) and prior.2 (0.980). We only considered genome-wide significant QTL SNPs within +/-250kb of the GWAS lead SNP of each locus. Phased LD matrices from 1000 Genomes, phase 3 (EUR), and sample overlap correction were used for the colocalization analysis. We started with 2-traits analyses comparing GWAS and each QTL one at a time: GWAS-eQTL, GWAS-meQTL, and GWAS-sQTL. Then, we performed 3-trait (G-e-m, G-s-e, G-s-m) and 4-trait (G-e-m-s) analyses. For each matrix (trait x SNP), one gene/probe per trait is selected at a time. Any matrix (trait x SNP) from 2,3,4-trait analyses is dropped if there are less than 50 SNPs. The colocalization events showing the consistent number of tested traits and colocalizing traits were included as the final result. For sensitivity analysis, we performed a similar multi-QTL colocalization with the stricter prior.2 parameter in HyPrColoc: 0.99 and 0.995.

### Imputed methylome-wide association study

We performed an imputed methylome-wide association study (MWAS) by predicting the function/molecular phenotypes into GWAS using the same melanoma meta-analysis as for the multi-QTL colocalisation^4^ and methylation data from both TCGA SKCM and melanocyte data. TWAS FUSION^35^, which was originally designed for transcriptome-wide association studies (TWAS), was adapted to perform the MWAS analysis. To summarize, we first collected the summary statistics without any significance thresholding. We then computed functional weights from our melanocyte methylation data one CpG Probe at a time. Probes that failed to pass a heritability check (minimum heritability *P*-value of 0.01) were excluded from the further analysis. A *cis*-locus was restricted to 50kb on either side of the CpG Probe boundary. For melanocyte data, from 386,520 probes meeting basic quality control, 21,252 probes passed the heritability check and were included as MWAS weights for association analysis using the melanoma GWAS summary stats and 1000 Genome, phase3 (EUR) LD reference. For the MWAS results, a genome-wide significance cutoff (MWAS *P*-value < 0.05/number of probes tested) was applied.

### eQTL/meQTL mediation analysis

A workflow (**Figure S9**) was applied to identify the potentially colocalized eQTL-meQTL pairs sharing a common causal variant followed by mediation and partial correlation analysis as originally described by Pierce and colleagues^17^. To identify candidate eQTL-meQTL pairs, we first restricted the meQTL analysis to lead SNPs (eSNPs) for each eGene from eQTL results and significant CpG probes (meProbes) from meQTL results to re-identified *cis*-meQTLs to these eSNPs. To reduce the redundant associations with the same SNP linking to a cluster of CpGs, we pruned our list of CpG probes by keeping only the CpGs whose lead meSNP had the highest LD with a lead eSNP. As a result, we identified each eGene paired with only one meCpG (eGene-meCpG pair), whose lead meSNP was in the strongest LD with the eSNP. In our melanocyte data, there were a total of 2,374 eGene-meCpG pairs showing association with a common SNP and available for colocalization analysis. HyPrColoc^34^ was used to perform colocalization analysis with the parameter prior.1 = 1e-4. A total of 841 potentially “colocalized” eQTL-meQTL pairs (including 296 common SNPs) were selected for downstream mediation analyses based on the posterior probability of a common causal variant (CCV) above 0.8.

For mediation analysis, we used our melanocyte data on 106 genotyped individuals with both expression and methylation data to conduct tests of mediation for two hypothesized pathways: (1) SNP -> Methylation -> Expression or “SME”, and (2) SNP -> Expression -> Methylation or “SEM”; For all lead eSNPs, the *cis*-eQTL association was re-tested with adjustment for methylation of the CpG (and vice versa). The difference between the beta coefficients before and after adjustment for the *cis* gene was expressed as the “proportion of the total effect that is mediated” (i.e., % mediation), calculated as |(βunadj – βadj)|/|βunadj|, with βunadj and βadj representing the total effect and the direct effect of the variant, respectively^17, 36^. All regression analyses were adjusted for PCs inferred from expression or methylation data. The Sobel *P*-value for mediation was calculated using the same formula as in previous publications^17, 37^.

We also performed the partial correlation analysis using the co-localized eQTL-meQTL pairs in our 106 melanocyte datasets. The Pearson correlation coefficients between the expression gene and the methylation probe were calculated after adjusting for expression and methylation PCs, respectively. Both the expression gene and methylation probe were regressed on the lead eSNP, and the residuals from these regressions were obtained as the expression and methylation values that lack the phenotypic variance due to the effect of the SNP. Correlation coefficients before and after SNP adjustment were compared to identify the eGene-meCpG pairs showing the partial correlation. To explore the extent to which partial correlation could be due to secondary, co-localized causal variants affecting both the expression trait and the CpG being analyzed, we also searched for secondary association signals for the eGene-meCpG pairs with partial correlation *P* < 0.05 and colocalization CCV > 0.8. For 73 pairs meeting these criteria, we adjusted for both the primary and secondary lead eSNP-meSNP. After this adjustment, 63 pairs were still significant (P > 0.05) (**Table S9**).

To explore the potential influence of CpG probes exclusion on methylation-expression mediation analysis, we pulled the 5,575 methylation probes that were dropped from melanocyte meQTL analysis. These are probes with SNPs of MAF>0.05 in EUR (minfi function dropLociWithSnps using SNPs parameters: “SBE” and “CpG”) that were excluded to avoid technical artifacts affecting genotype effect on allelic methylation levels, as suggested by other studies^12, 38^. Among them, 583 unique methylation probes overlapped (within +/-1bp) with 594 unique melanocyte eQTL SNPs (595 unique probe-SNP pairs and 925 unique probe-SNP-gene trios). When overlaid with melanoma GWAS-melanocyte eQTL colocalization results (using HyPrColoc), none of the 594 eQTL SNPs overlapped with melanoma GWAS colocalized SNPs (posterior probability>0.8) or their proxies (r^2^ > 0.8). Ten of the 594 eQTL SNPs were the strongest eQTL SNPs of an eGene (eSNP). Predicted allelic transcription factor binding for these ten SNPs was searched on Haploreg v4.1 (https://pubs.broadinstitute.org/mammals/haploreg/haploreg.php).

### Identifying *cis*-mediators for *trans*-meQTLs

To explore the mediation of *trans*-meQTL by *cis*-eQTL (e.g. of potential transcription factors), mediation analysis was performed by applying eQTLMAPT^39^ to the primary melanocytes’ meQTL data. Only trios with evidence of both *cis*-eQTL and *trans*-meQTL association were included. To detect the mediation effects, 152 candidate trios were derived from significant *cis*-eQTL and *trans*-meQTL associations (based on *FDR* < 0.05 and < 0.01, respectively). We performed the mediation analysis with an adaptive permutation scheme and GPD approximation with parameters N = 10,000 and α = 0.05 for all candidate trios. All PEER factors included in eQTL and meQTL analyses and other covariates (top 3 genotype PCs) were adjusted and trios with suggestive mediation were reported using mediation *P*-value threshold < 0.05.

### Enrichment of melanoma GWAS variants in meQTLs

We generated quantile-quantile (QQ) plots to evaluate whether melanoma GWAS variants were enriched in meQTLs of melanocyte or TCGA SKCM. To minimize the impact of linkage disequilibrium (LD) on the enrichment analysis, we performed LD-pruning to identify independent SNPs among all the GWAS variants using PLINK v1.90 beta^40^ (r^2^ = 0.1 and window size 500 kb). QQ plots were made using *P*-values (-log10) of melanoma GWAS^4^ for non-meQTL SNPs v.s. meQTL SNPs after LD-pruning.Deviation from the 45-degree line indicates that melanoma GWAS SNPs are enriched in meQTL SNPs.

### Functional annotation of CpGs and meQTLs

Functional annotation of CpGs and meQTLs has been described previously^12^. We annotated 10 genomic features of CpGs, including CpGs located in CpG Islands, low or high CpG regions, promoters, enhancers, gene bodies, 3 prime untranslated regions (3’UTR), 5’UTR, 0–200 bases upstream of transcription start sites (TSS200), and 201–1500 bases upstream of transcription start sites (TSS1500). Hypergeometric tests were used to evaluate if the identified *cis-* and *trans-*meQTL CpGs showed enrichment for CpGs annotated with those genomic features. The significance threshold was defined by a fold change of >1.2 or < 0.8 and a Bonferroni-corrected threshold *P* < 0.05/10=0.005.

In addition, we determined the distribution of genome-wide meCpG probes based on their genomic position in relation to CpG islands and nearby genes. Enrichment fold change was calculated as the ratio of the fraction of meQTLs overlapping with genomic annotations v.s. the fraction of randomly selected SNPs overlapping with the genomic annotations. ‘epitools’ was used for this analysis.

### Motif enrichment analysis for *trans*-meQTL

Enrichment of known sequence motifs among trans-CpGs was assessed using the PWMEnrich package in R (https://bioconductor.org/packages/PWMEnrich/). One hundred and thirty-one CpG probes with *trans*-meQTL association with rs12203592 were selected for enrichment analysis. For PWMEnrich, the 101-bp sequence around each interrogated CpG site was used, and unique 2kb promoters in humans were used as the pre-compiled background set.

### IRF4 ChIP-sequencing in melanoma cells

To identify genome-wide binding sites of IRF4 in melanoma cells, we performed ChIP-sequencing against eGFP tagged IRF4. We generated an inducible eGFP tagged IRF4 cell line in 501Mel cells by cloning eGFP tagged IRF4 downstream of the tetracycline response element in a PiggyBac transposon system^41^. We used the Tetracycline-ON system where the expression of eGFP-IRF4 can be induced by adding doxycycline or tetracycline. For ChIP experiments, the eGFP-tagged IRF4 expressing 501Mel cells were grown on ten 10 cm dishes, and 1 ug/ml of doxycycline was added for the induction. Chromatin immunoprecipitation was performed according to Palomero and colleagues^42^ as follows: Twenty million cells were crosslinked with 0.4% formaldehyde for 10 minutes at room temperature, quenched by 0.125M glycine for 5 min at RT and chromatin was then sheared by 5 min sonication (25% amplitude, 30sec off and 30 sec on) using a probe sonicator (Epishear, Active Motif). Immunoprecipitation was performed with Protein G Dynabeads (Life Technologies), with a total of 10 µg of anti-GFP antibody (3E6 from Molecular Probes, #A-11120). The bead-bound immune complexes were washed five times with wash buffer (50M Hepes pH 7.6, 1mM EDTA, 0.7% Na-DOC, 1% NP-40, and 0.5M LiCl) and once with TE. Crosslinking was reversed by washing the immune-complexes and sonicated lysate input in elution buffer (50mM Tris pH 8, 10mM EDTA, 1% SDS) overnight at 65 °C. Then the samples were treated with 0.2 μg/μL of RNase A for 1 hour at 37°C followed by treatment with 0.2 μg/μL proteinase K for 2 hours at 55°C. DNA was extracted from the samples using phenol:chloroform. ChIP-seq DNA libraries were prepared from the purified ChIP DNA and input DNA using the NEBNext ChIP-seq Library Prep Kit (E6200, NEB). Libraries were prepared from 8-15 ng of fragmented ChIP or input DNA, which were amplified with 10 PCR cycles. The amplified libraries were purified using Agencourt AMPure XP beads (A63881, Beckman Coulter) and then were paired-end sequenced. Approximately 30 million raw reads were mapped of each sample to the human hg19 reference genome using Bowtie 2^43^. The aligned reads were then used as an input for peak calling using MACS^44^.

### IRF4 knockdown and RNA sequencing

The human melanoma cell line 501mel was cultured in RPMI-1640 cell culture medium (Gibco) supplemented with 10% FBS (Gibco) in a humid incubator at 5% CO2 and 37°C. IRF4 was knocked down in three biological replicates of 501mel cells by transfecting the cells using Lipofectamine (RNAiMAX, Thermo Fisher) with siRNA (Silencer Select #AM16708, Thermo Fisher) for 48 hours. Cells were harvested and RNA was extracted using Quick-RNA Mini prep (#R1055, ZYMO Research). IRF4 knockdown was verified by RT-qPCR before generating sequencing libraries. RNA-sequencing was performed on the NovaSeq 6000 system and ∼150 million raw reads were mapped to human transcriptome GRCh38 using Kallisto^45^ and differential expression analysis was performed using Sleuth^46^.

## Results

### Identification of cell-type-specific melanocyte meQTLs

To establish a melanocyte-specific meQTL dataset, we assessed DNA methylation levels in cultured melanocytes from 106 newborn males mainly of European descent using Illumina 450K methylation arrays (**Material and Methods**). We then performed *cis*-meQTL analysis assessing variants within +/-1Mb of each CpG probe and identified 13,274 unique CpG probes (meProbes) with 1,497,502 significant *cis*-meQTLs before LD-pruning (**Table S1A**). Most *cis*-meQTL variants are clustered near CpGs (< ∼100kb), where variants closer to the target CpGs tended to have lower *P*-values and larger effect sizes (**Figure S1**). Among 13,274 meProbes, 29% were located in CpG islands and 34% in CpG-adjacent regions (shores and shelves), with the rest (38%) away from CpG islands (Open Seas) (**Figure S2**). meProbes are also mainly located in or near the gene body (73% are within 1500bp of Transcription Start Sites, UTRs, 1st exon, or gene body), and the rest (27%) in intergenic regions. Compared to non-meProbes, meProbes are most enriched in Open Seas and intergenic regions, while most depleted in islands and 1st exons. At the variant level, *cis*-meQTLs are also significantly depleted in CpG islands and gene promoter regions (**Figure S3**).

To supplement these meQTLs, as well as melanocyte-specific eQTLs we previously identified ^6^, we also performed mRNA splice junction QTL (sQTL) analysis using previously generated RNAseq data from the same melanocytes through which we identified 7,054 unique splice junctions with 887,233 *cis*-sQTLs (not LD-pruned) (**Table S1A**). Together with our previous eQTL findings, we identified a total of 1,039,047 non-overlapping eQTL/meQTL/sQTL variants in melanocytes, a substantial proportion (40.4%) of which are only detected by meQTL (**Figure S4**). Of meQTL variants, 27.4% and 21.8% were also detected as eQTL and sQTL, respectively, and 13.3% were significant for all three QTLs. Among eQTL variants, 42.3% and 36.7% were also detected as meQTL and sQTL, respectively. Among sQTLs, 44.2% and 40.5% displayed an overlap with eQTLs and meQTLs, respectively.

### Multi-QTL colocalization improved melanoma GWAS annotation

To explore the contribution of cell-type specific meQTL and other QTLs to melanoma GWAS annotation, we first performed multi-trait colocalization using HyPrColoc^34^ using summary data from a recent melanoma GWAS meta-analysis of 36,760 histologically confirmed and self-reported cases^4^. Melanocyte meQTLs colocalized with melanoma GWAS signals (posterior probability > 0.8) at 13 of 54 loci, while sQTLs displayed colocalization at two loci (**Figure 1**, **Table S1B, S2**). Together, at least one of three QTL types colocalized with melanoma GWAS signal at 21 of 54 melanoma loci (39%), which is a considerable improvement from the 12 loci (22%) explained by eQTLs alone using the same approach (HyPrColoc; note that this percentage differs slightly from 16% reported in Landi *et al*^4^, where eCAVIAR^47^ was used for colocalization). Sensitivity analysis, adjusting the second prior from 0.98 to 0.99 and 0.995, indicated that 80% (61/76) and 64% (49/76) of colocalization events were still detected for the same traits at posterior probability > 0.8, respectively (**Table S2**). These data demonstrated that cell-type-specific multi-QTL colocalization could explain close to half of melanoma GWAS loci and that methylation QTL is the largest contributor colocalizing with 24% of the known loci.

**Figure 1.**
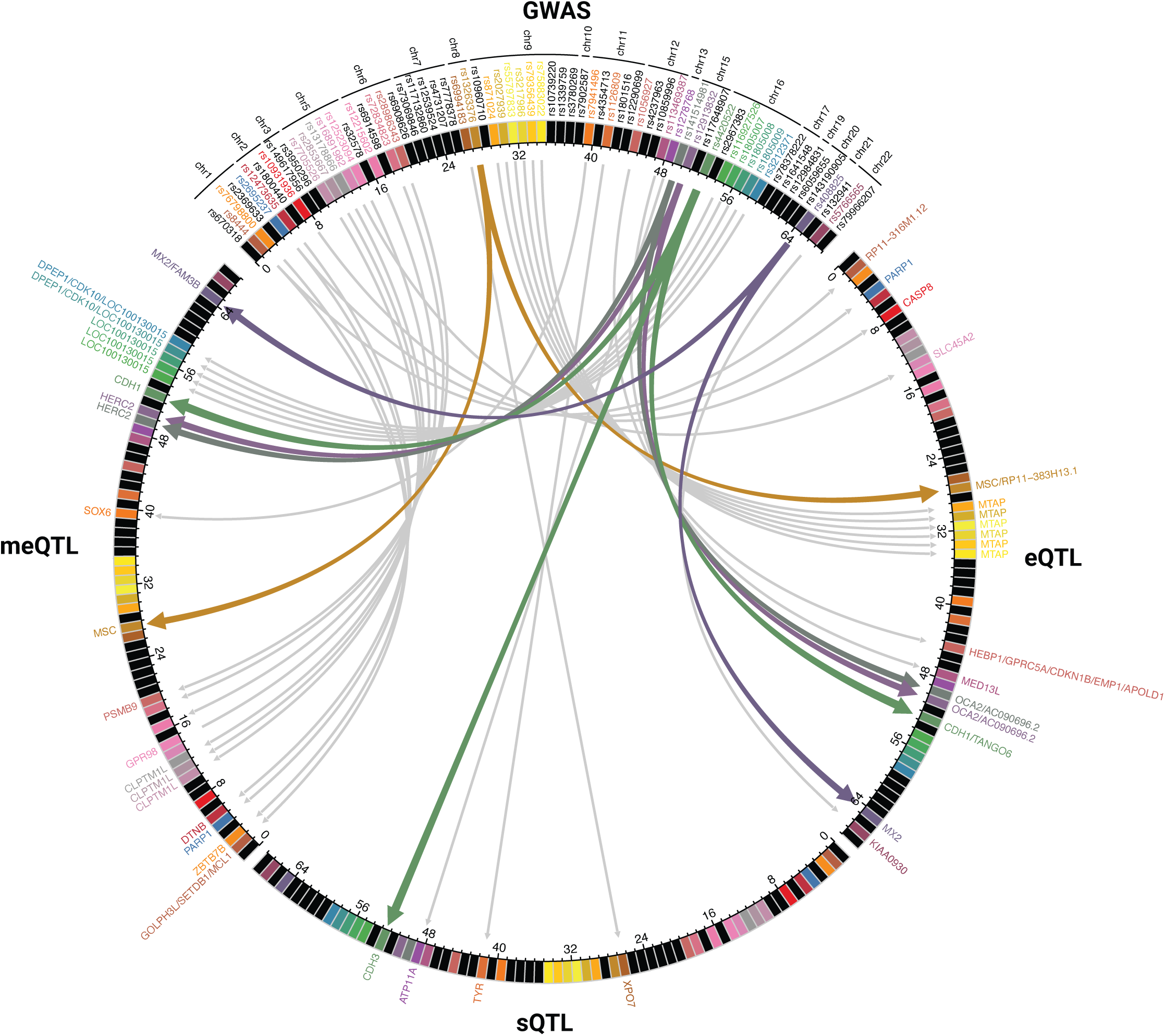
Melanocyte meQTL and multi-QTL colocalization improved melanoma GWAS annotation. Circos plot shows significant colocalization of melanoma GWAS loci (top) with eQTLs (right), sQTLs (bottom), and meQTLs (left). Colocalization between individual GWAS loci with multiple QTL traits are depicted by thicker, colored lines. GWAS loci are sorted by genomic coordinate and labeled with GWAS Lead SNPs with different colors; GWAS loci without any colocalizing-QTL are shown in black. QTL-associated gene symbols are also labeled with the same color as GWAS loci.

Further, multi-QTL colocalization identified four loci where more than one cell-type-specific QTL trait colocalizes with the melanoma GWAS signal (**Figure 1**, **Table S2**). For three loci, both eQTL and meQTL colocalized with the GWAS signal (*MSC/RP11-383H13.1, OCA2/AC090696.2,* and *MX2*; **Figure S5A-C**). At the fourth locus, all three QTL traits, including eQTL (*CDH1*), meQTL (meCpG near *CDH1*), and sQTL (splice junction in *CDH3* gene), were colocalized with the GWAS signal (**Figure S5D**). For the locus near *MX2*, colocalization identified rs398206 as a common causal variant for eQTL, meQTL, and melanoma risk, validating our previous findings identifying this variant as a functional *cis*-regulatory variant regulating *MX2*^48^. Here, meQTLs for two CpG probes in the gene body display the same allelic direction of effect as that of *MX2* eQTL, where higher methylation levels are correlated with the allele associated with increased *MX2* expression, consistent with the observations that DNA methylation in the gene body is positively correlated with gene expression. *OCA2* is a known pigmentation gene and, within this locus, the lead GWAS SNP located in the *HERC2* gene, rs12913832, was identified as a common causal variant for eQTL, meQTL, and melanoma risk through the expression of both *OCA2* and an antisense *HERC2* transcript, *AC090696.2.* These results are consistent with the previous findings that a melanocyte-specific enhancer encompassing rs12913832 regulates *OCA2* expression through an allele-preferential long-range chromatin interaction^49^. The *MSC/RP11-383H13.1* locus was initially identified as a novel locus by melanoma TWAS using our melanocyte eQTL dataset^6^ and data from a prior melanoma GWAS meta-analysis^5^, with this locus being subsequently identified as a genome-wide significant GWAS locus by the larger melanoma GWAS^4^. Our multi-QTL colocalization indicated that DNA methylation is also involved in this locus in mediating melanoma risk. Finally, for the *CDH1/3* locus, rs4420522 in the intron of *CDH1* was identified as a common likely causal variant for an eQTL (*CDH1*), meQTL (*CDH1* gene body open sea CpG), sQTL (*CDH3*), and melanoma risk. Notably, the eQTL (*CDH1*) and sQTL (*CDH3*) are for two different homologous genes encoding E-cadherin and P-cadherin, respectively, that are located adjacent to each other. Same variants being an eQTL for one gene and sQTL for another has been shown for a subset of GTEx sQTLs in a recent study^50^, but whether they share candidate causal variants was not clear. Here we show an example of a common candidate causal variant affecting gene expression or splicing of two different genes in the same cell type.

### Imputed MWAS identified novel melanoma-associated loci

Given that meQTLs colocalize for a sizable proportion of melanoma GWAS signals, we further performed an imputed methylome-wide association study (MWAS)^26^ using the melanocyte methylation data. Adopting the approach used for transcriptome-wide association studies (TWAS)^35^, we trained models of genetically regulated CpG methylation in our melanocyte dataset (**Material and Methods**) and tested the association of imputed methylation levels and melanoma risk using the summary statistics from melanoma GWAS. Significant MWAS was observed for 159 meCpGs (Bonferroni-corrected MWAS *P* < 0.05/21,252 tested probes), which overlapped 29 known genome-wide significant melanoma GWAS loci and further nominated 10 potentially “new” loci (**Table S3A, S4**). Among these new loci, six overlapped with GWAS loci previously identified in a pleiotropic analysis between melanoma and nevus count and/or melanoma and hair color traits, or loci identified by melanocyte TWAS (*NIPAL3, NOTCH2, HDAC4, AKAP12, CBWD*, and *SYNE2*; **Table S4**)^4^, suggesting that the MWAS approach effectively identifies bona fide susceptibility loci found via complementary approaches. Besides these six loci, the other four loci included CpG probes on or near *SPOPL, NUMA1/LRTOMT, SNORD41/TNPO2, EPB41L1*, and *RPRD1B*. These results demonstrated the potential of MWAS to nominate candidate susceptibility genes that are missed in the single-variant analysis.

Consistent with our comparisons between eQTL and meQTL colocalization, MWAS and TWAS together explained 54% of melanoma GWAS loci, which is a considerable improvement from 28% of GWAS loci by TWAS alone (**Table S3A**)^4^. Combined with the findings from colocalization analyses, melanocyte eQTL and meQTL together explained 63% of melanoma GWAS loci (**Table S3B**). TWAS, MWAS, and multi-QTL colocalization cross-validated each other in 18/54 (33%) of GWAS loci, where one or more approaches pointed to the same affected genes (**Figure 2**). Of the 16 genes that were supported by both TWAS and MWAS (gene assignment is based on CpG probes within 1.5kb of TSS, 5’ UTR, 1st exon, gene body, or 3’ UTR of a gene), 6 genes displayed the same direction of effect relative to melanoma risk (Z-scores in the same direction), while 5 genes displayed the opposite direction of effect (**Table S4**). However, the other 5 genes were matched with CpG probes displaying the effect in both directions. These data suggest potential co-regulation of gene expression and promoter CpG methylation in these loci, contributing to melanoma risk.

**Figure 2.**
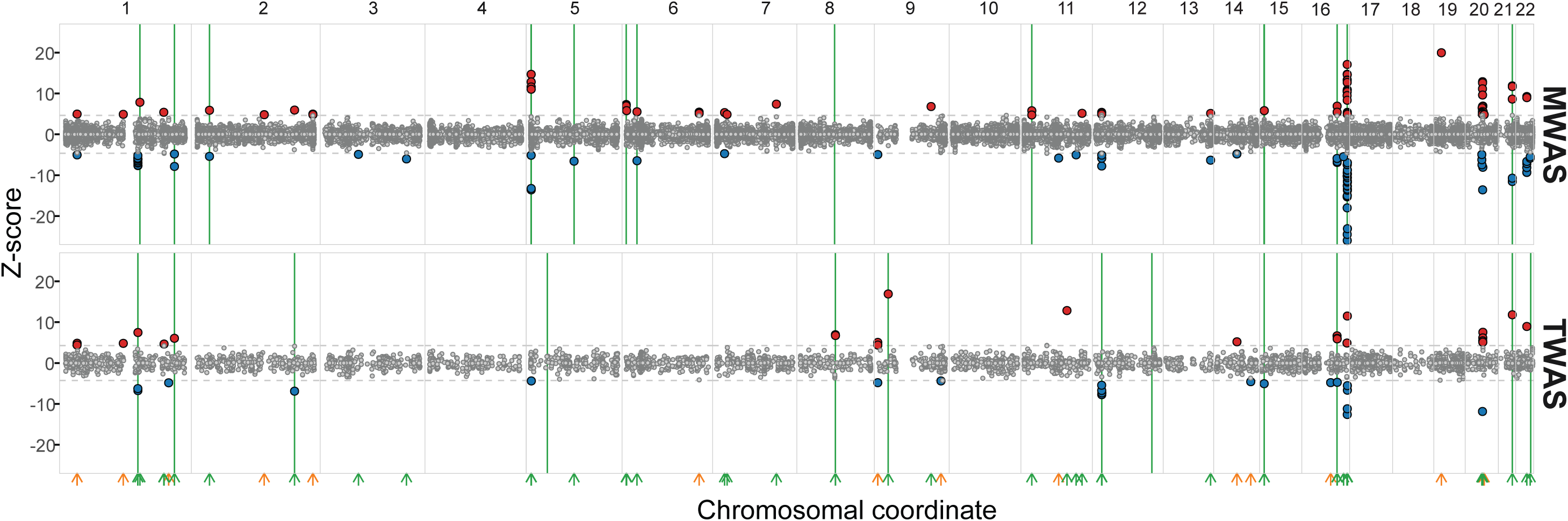
Manhattan plots of melanocyte TWAS and MWAS results combined with findings from eQTL and meQTL colocalization. Each circle represents the TWAS or MWAS z-score of a gene (TWAS) or a CpG probe (MWAS) reflecting significance and the direction of effect relative to melanoma-risk (red: higher-level correlates with melanoma risk, blue: lower-level correlates with melanoma risk). Z-scores are shown on the y-axis, and chromosomal positions are on the x-axis. Green arrows: overlapping melanoma GWAS loci, orange arrows: new loci detected by TWAS or MWAS, green lines: colocalization of eQTL or meQTLs with melanoma GWAS loci. Gray dashed horizontal lines: significance threshold defined by 0.05/number of probes or genes tested.

Through both colocalization and TWAS/MWAS, melanocyte eQTL and meQTL nominated a total of 107 unique candidate melanoma susceptibility genes. Ingenuity Pathway Analysis (http://www.ingenuity.com/index.html) identified biological pathways enriched by these genes including those in melanin biosynthesis (L-dopachrome biosynthesis, L-DOPA degradation, eumelanin biosynthesis), apoptosis (apoptosis signaling, Myc-mediated apoptosis signaling, retinoic acid-mediated apoptosis signaling), autophagy, adhesion junction signaling (epithelial adherens junction signaling, remodeling of epithelial adherens junctions), and melanoma-specific signaling (melanoma signaling, Wnt/beta-catenin signaling), among others (**Table S5A**). Of these, melanoma-specific signaling and apoptosis pathways are strengthened by adding meQTL compared to a similar analysis using only eQTL in melanocytes and skin tissues^4^. Notably, upstream regulator analysis identified the transcription factor MITF as the most significant regulator of these genes (**Table S5B**), which is consistent with its known role as the master regulator of melanocyte lineage^51^ and a melanoma susceptibility gene^52, 53^. Together, these data demonstrated that meQTL from the cell-type of disease origin is complementary to eQTL data and greatly increases the power to nominate candidate causal genes.

### Melanocyte meQTLs are substantially preserved in melanomas

Given the large contribution of melanocyte meQTLs underlying melanoma GWAS loci, we further asked if and how well the genetic control of CpG methylation in the melanocytic lineage is preserved in malignant melanomas. For this, we performed a meQTL analysis of 444 cutaneous melanomas from TCGA using data generated from the same 450K methylation array platform and using the same analytic approach, except for adding regional genomic copy number as a covariate (**Material and Methods**). First, we identified 3,794,446 genome-wide significant *cis*-meQTLs (not LD-pruned) for 15,308 unique meProbes from TCGA melanomas, which are higher numbers than those observed from melanocytes (15% more meProbes). When meProbes were compared between datasets, 45.9% of melanocyte meProbes were also significant in melanomas, while 39.8% of melanoma meProbes were observed in melanocytes (**Figure S6A**). Melanocyte meQTL preservation in melanoma is even higher at the gene level, showing 65% preservation when meProbes are assigned to genes based on their position relative to gene bodies or promoters (**Figure S6B**). The effect sizes of the best meQTL for each meProbe were highly correlated for 6,087 common meProbes in both groups (*P*-value < 2.2e-16; R = 0.74), with 88.4% of them displaying the same direction of effect (**Figure S7**). We further calculated the true positive rates (π1) of top *cis*-meQTLs (*FDR* < 0.05) from melanocytes by examining their *P*-value distributions in melanoma meQTLs, and vice versa. The true positive rate (π1) was 0.825 and 0.822 for melanocyte meQTLs in melanomas and melanoma meQTLs in melanocytes, respectively, displaying a high level of meQTL preservation between two datasets. Notably, normal to tumor preservation was much less at the eGene level, where only 12.7% of melanocyte eQTL genes were preserved in melanomas (**Figure S6A**), in contrast to the high preservation rate of meQTL. Among 635 preserved eGenes, 230 (36%) were associated with one or more preserved eProbes.

We then investigated whether melanoma-specific meQTL corroborates melanoma GWAS annotation through colocalization and MWAS. Melanoma meQTLs colocalized with the melanoma GWAS signal at 11 of 54 loci (20%) (**Table S6**), and melanoma MWAS overlapped with 19 GWAS loci (35%) and further identified six new loci (**Table S7**). Among these were loci only explained by melanoma meQTL but not by melanocyte QTLs; melanoma meQTLs uniquely annotated five GWAS loci (CpG probes near *PPARGC1B*, *OBFC1*, and *SHANK3*) and identified four novel MWAS loci (**Figure 3**; **Table S8**). Through colocalization and MWAS, melanoma meQTL explained 46% (25/54) of melanoma GWAS loci, which, despite the >4-times larger sample size and an overall higher number of identified meProbes, is considerably less than that by melanocyte meQTL (56%). Consistent with this observation, melanoma risk-associated variants are more enriched for melanocyte meQTLs than for melanoma meQTLs (**Figure S8**). Thus, these data demonstrate that genetically regulated CpG methylation observed in the melanocyte-lineage is substantially preserved in tumors. Nevertheless, these data also show that cancer susceptibility reflected in GWAS signals is better explained by DNA methylation from normal homogeneous cells of disease origin than by that from heterogeneous tumor tissues, even with considerably larger sample size. Overall, melanocyte multi-QTL and melanoma meQTL collectively explain 39 melanoma risk-associated loci, representing 72% of all known genome-wide significant loci.

**Figure 3.**
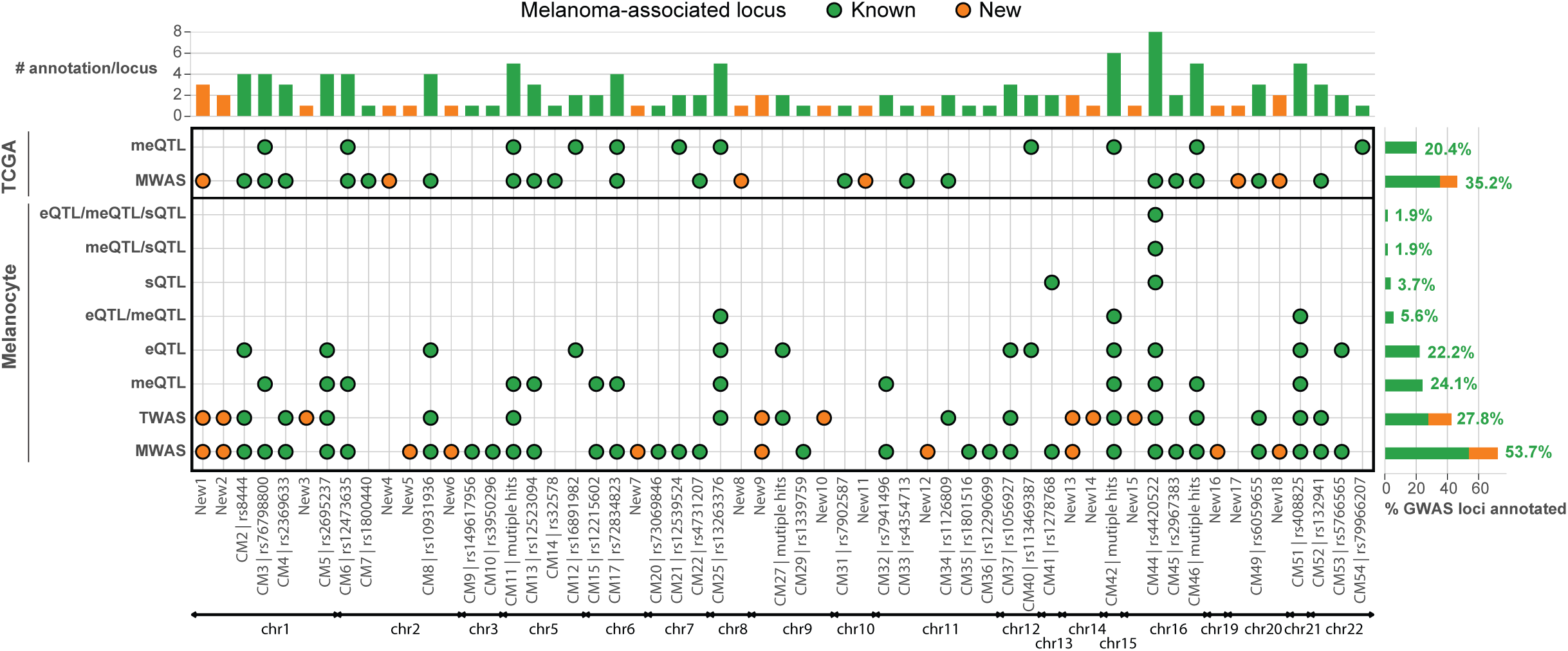
Summary of melanoma GWAS annotation using melanocyte multi-QTL and TCGA-melanoma meQTL. Known melanoma-associated loci (green circles) are defined by the findings from the newest melanoma meta-analysis. The new melanoma-associated loci (orange circles) are identified based on TWAS or MWAS analysis. Known and new GWAS loci are sorted by genomic coordinate. The top boxplot shows the total number of annotations per locus by multi-QTL colocalization (shown by QTL types) or TWAS/MWAS from melanocyte and TCGA datasets. The right marginal boxplot shows the percentage of GWAS loci annotated by each approach (the percentage of the known loci is labeled in green).

### Genetic control of DNA methylation and gene expression in melanocytes

To investigate the genetic control of gene expression and DNA methylation in primary melanocytes beyond their contribution to melanoma risk, we sought to determine whether eQTLs and meQTLs more broadly share the same causal variants and whether one has a causal effect on the other. For this, we performed colocalization of eQTLs and meQTLs followed by mediation and partial correlation analysis as previously described by Pierce and colleagues ^17^ (**Figure S9**). We first took 4,886 unique eSNPs (strongest eQTL SNP for each eGene) from eQTL data and re-identified *cis*-meQTLs, limiting to these 4,886 SNPs and 13,274 meCpG probes (meProbes). After pruning overlapping meProbes, we identified 2,374 unique eGene-meProbe pairs linked by the same eSNP, 841 of which (35%) were colocalized at posterior probability > 0.8 using HyPrColoc (prior1 = 1e-4; prior2 = 0.95; **Material and Methods**). We then performed partial correlation analysis for those 841 eGene-meProbe pairs, of which 50 (6%) displayed significant partial correlation after conditioning on the primary and the secondary independent variants (*FDR* < 0.05), and 197 (23%) displayed correlation at a relaxed cutoff when conditioning on the primary variant (*P* < 0.05) and 187 when conditioning on both the primary and the secondary variants. These data suggested a link between DNA methylation and gene expression beyond that through common causal variants (**Figure S10A**; **Table S9; Methods**). Next, we performed mediation analysis for 841 eGene-meProbe pairs to estimate the effect of SNP on gene expression mediated by DNA methylation and vice versa. The results indicated that 32 unique eGene-meProbe pairs (4%) displayed significant mediation either of methylation on expression (25 pairs; *FDR* < 0.05 & % mediation >0) or of expression on methylation (25 pairs; *FDR* < 0.05 & % mediation > 0), where 18 pairs were significant under both hypotheses (**Figure 4**; **Table S10**). All 32 significantly mediated pairs were included in 197 pairs displaying a marginal partial correlation (*P* < 0.05) (**Figure S10B**). Among 197 SNP-gene-probe trios, 69% (135 trios) displayed an opposite allelic direction of effect between meQTL and eQTL, while 31% (62 trios) displayed the same allelic direction of effect.

**Figure 4.**
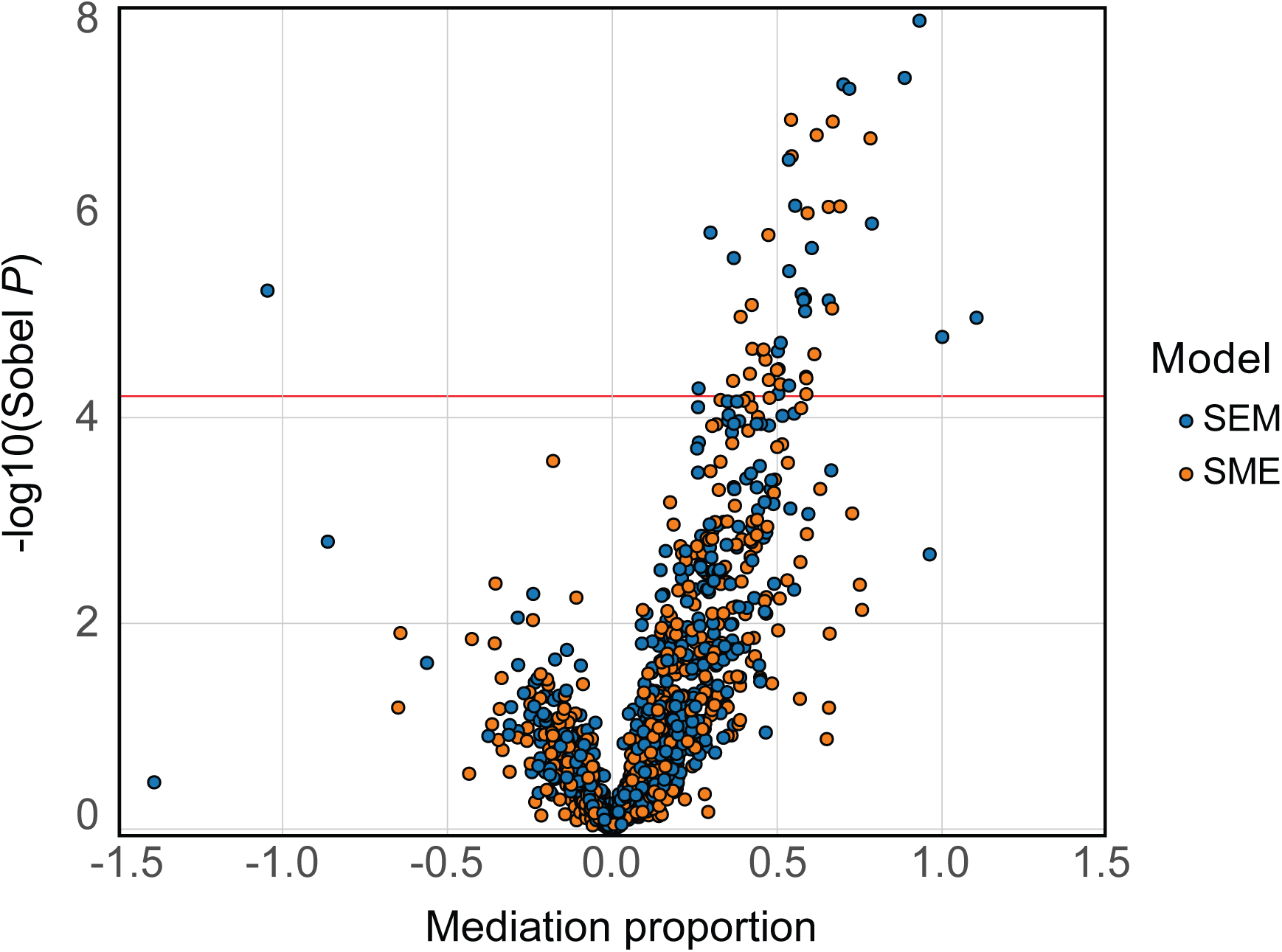
Mediation analysis of potentially colocalizing SNP-eGene-meProbe. The volcano plot shows the mediation analysis results for both the SEM (blue) and SME (orange) models. Sobel P indicates the significance of the mediation analyses, where the red horizontal line indicates *FDR* = 0.05 cutoff. The mediation proportion shows the proportion of the total effect (*cis*-meQTL) mediated by a *cis*-Gene (SEM) or the proportion of the total effect (*cis*-eQTL) mediated by *cis*-Probes (SME). Mediation proportion can go in either direction, depending on the directions of the effects of the confounders with the *cis-*mediator, the confounder on the *cis*-gene or *cis*-Probes, and the non-reference allele on the *cis*-Probes or *cis*-Gene.

Our data suggested that a considerable proportion (∼35%; 841 of 2374) of eGene-meProbe pairs may share a causal variant between eQTL and meQTL in melanocytes. A subset (up to 23%; 197 of 841) of those displayed some evidence of methylation/expression co-regulation, where a majority displays an opposite directional effect. Ingenuity Pathway Analysis of 197 genes displaying significant mediation (*FDR* < 0.05) or partial correlation (*P* < 0.05) between melanocyte meQTL and eQTL highlighted pathways involving immune response and UVA-Induced MAPK Signaling among others (**Table S11**). Notably, 841 potentially colocalizing eGene-meProbe pairs were significantly enriched in melanocyte eGenes that are preserved in malignant melanomas compared to non-preserved eGenes (Fisher’s exact, *P* = 9.44e-07; OR = 1.68). We do not observe the same type of enrichment in preserved meProbes compared to non-preserved meProbes (*P* = 0.608; OR = 1.04). However, colocalizing eGene-meProbe pairs are significantly enriched in genes on or near the preserved meProbes compared to those with non-preserved meProbes (*P* = 3.36e-05; OR = 1.58). These data suggest that genetic influence on potentially co-regulated DNA methylation and gene expression in primary melanocytes tend to be well maintained during malignant transformation.

Although conventional meQTL analyses using array-based methylation measurement exclude SNPs overlapping CpGs themselves, SNPs on CpG sites could potentially have a high impact on allelic methylation and target gene expression. Among all the SNPs in CpG, 10.6% were significant eQTLs in melanocytes. Of these, we focused on 10 CpG SNPs that are the strongest eQTL for an eGene (eSNPs) in our melanocyte dataset (**Table S12**). A majority of these CpG probes were located in promoter or enhancer regions near TSS. While the allelic changes from C or G to A or T are considered to abolish the CpG sites preventing methylation, some of them were predicted to create transcription factor binding sites in exchange. In an example of cg16139068, a CpG probe near the TSS of *OGDHL,* an allelic change from CpG to CpA (rs61846889) dramatically increases predicted binding affinity for Ahr::Arnt::HIF1 complex (Haploreg v4.1 position weight matrix from <u>doi:10.1093/nar/gkt1249</u>). rs61846889 is also a significant eQTL for *OGDHL* across multiple tissues as well as melanocytes, including sun-exposed skin (*P* = 1.2e-20, normalized effect size relative to A allele = 0.62; GTEx v8). These data hint at a hypothesis that CpG SNPs could lead to allelic gene expression by directly affecting DNA methylation while simultaneously affecting transcription factor binding. Together our data provide insights into an intersection of eQTL and meQTL in genetic control of gene expression and DNA methylation in melanocyte biology.

### Melanocyte *trans*-meQTLs highlight an IRF4 transcriptional regulatory network

Next, we performed *trans*-meQTL analysis of melanocytes, testing SNPs outside the +/-5Mb boundary of each CpG probe or on a different chromosome. We observed 332 unique CpG probes with one or more significant *trans*-meQTL at *FDR* < 0.01 (**Table S13; Figure 5**). For 65% (215 of 332) of those CpG probes, the best *trans*-meQTL variant was also a significant *cis*-eQTL in melanocytes. Among all the significant *trans*-meQTL variants, only one variant was a hot spot *trans*-meQTL for more than 10 CpGs across the genome. Namely, rs12203592, a *cis*-eQTL for *IRF4* gene expression^6^, was a *trans*-meQTL for 131 CpGs (40%). rs12203592 was previously shown as a functional variant in melanocyte-lineage that regulates the expression of the IRF4 transcription factor^54^. In our previous study of melanocyte eQTLs, we identified rs12203592 as a significant *cis*-eQTL of *IRF4* as well as a genome-wide significant *trans*-eQTL for four different genes, *TMEM140*, *MIR3681HG*, *PLA1A*, and *NEO1*^6^, a subset of which displayed significant mediation by *IRF4 cis*-eQTL. In the current study, rs12203592 was identified as a *trans*-meQTL for two CpG probes (cg14710552 and cg07972322) located in *TMEM140* and one CpG probe (cg04330122) located in *PLA1A*, consistent with our findings in *trans*-eQTL. Furthermore, 95.4% (125 of 131) of rs12203592 *trans*-meQTL-CpG pairs displayed a positive effect size relative to the alternative T allele, where lower *IRF4* expression level is associated with higher methylation levels at the target CpGs (**Table S14**). These results are similar to the observation in blood samples, where *trans*-meQTL hotspots displayed consistent allelic directions^11, 12^. Our findings are consistent with the hypothesis that altered expression of *IRF4* by the *cis*-eQTL SNP, rs12203592, affects allelic methylation changes of those CpGs on or near multiple downstream target genes in melanocytes.

**Figure 5.**
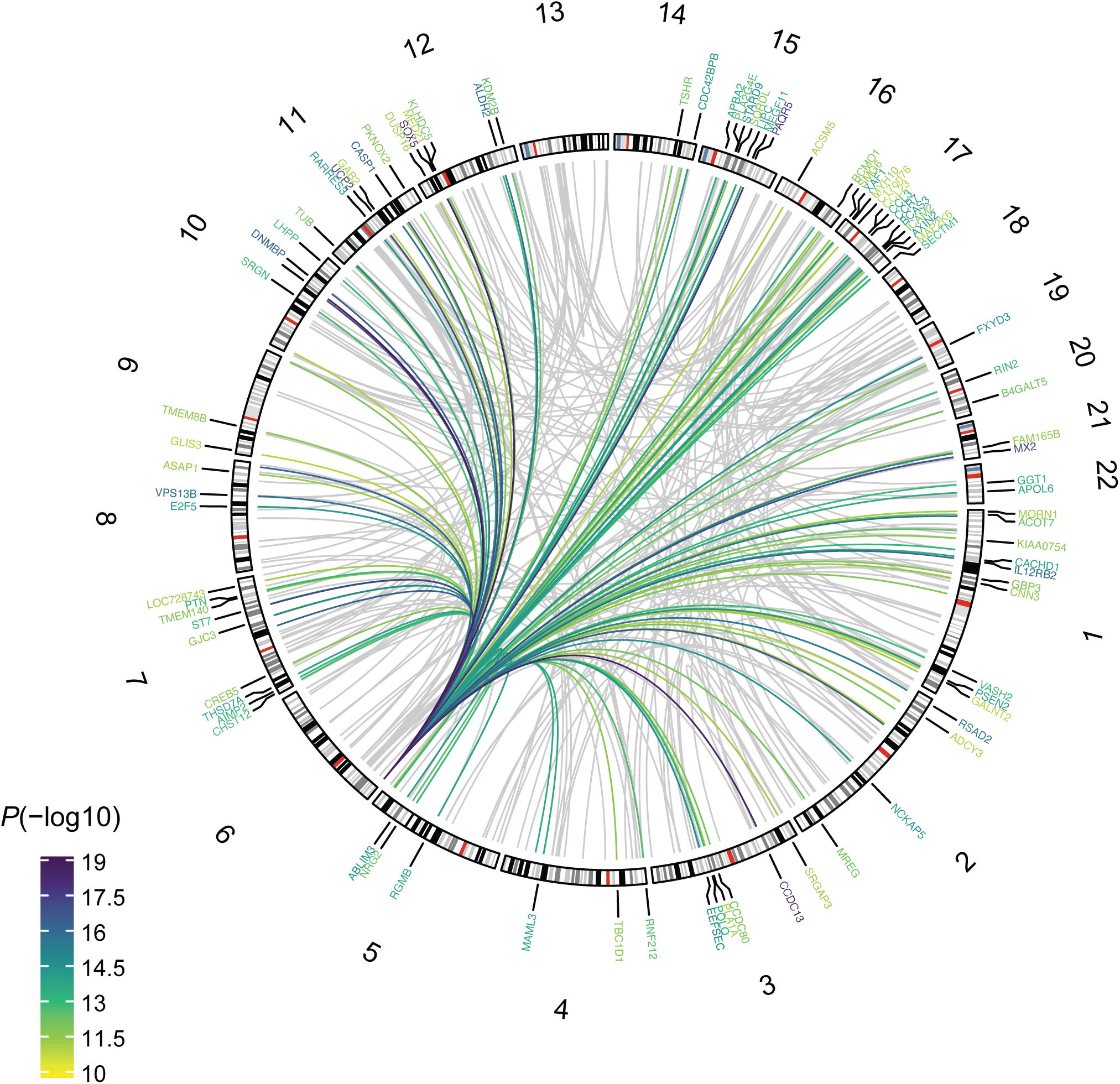
Melanocyte trans-meQTL. Circos plot shows the genome-wide significant *trans*-meQTLs at *FDR* < 0.01. The yellow-green gradient spikes show a hotspot *trans*-meQTL SNP, rs12203592, located at 6p25.3 that is associated with 131 CpG sites. Nearby genes of *trans*-meQTL associated CpG sites are labeled outside the circos plot.

We then asked if any *cis*-eQTL variant is driving *trans*-meQTL (i.e. via allelic expression of transcription factors and the subsequent effect on methylation of downstream targets) by performing mediation analysis using eQTLMAPT^39^. For this, we tested 152 *cis*-eQTL variant:*cis*-eQTL gene:*trans*-meQTL probe trios (*FDR* < 0.05 for *cis*-meQTL and < 0.01 for *trans*-meQTL), of which 24 trios displayed significant mediation at *P* < 0.05 (**Table S15; Figure S11**). An overwhelming majority of the significant trios (92%; 22 of 24) included rs12203592, where *cis*-eQTL of *IRF4* expression mediates the *trans*-meQTL effect of 18 putative target genes, further supporting IRF4-mediated target gene regulation in melanocytes. Notably, among 18 putative IRF4 target genes was a melanoma-risk associated gene, *MX2* (MX Dynamin Like GTPase 2)*. MX2* is an interferon-alpha-stimulated gene (ISG) with conventional roles in the innate immune response against HIV infection but was previously shown to have a melanocyte-lineage-specific function in promoting melanoma formation^48^. Similarly, IRF4 was originally known as one of the IFN-regulatory factors with roles in B and T lymphocytes^55–57^ but also has melanocyte-lineage specific roles in pigmentation traits^54^, which is consistent with its association with pigmentation traits^58, 59^, nevus counts^60^, and melanoma risk^4, 60^. These data suggest a melanocyte-specific functional interaction between two melanoma-risk associated genes, *IRF4* and *MX2*.

To further investigate if the targets of rs12203592 *trans*-meQTL are regulated by direct IRF4 binding, we performed IRF4 ChIP-seq using 501mel melanoma cells ectopically expressing *IRF4*. Among 131 significant *trans*-meQTL target CpGs (*FDR* < 0.01) of IRF4 *cis*-meQTL SNP rs12203592, 54 (41.2%) CpGs overlapped within +/-100bp of IRF4 ChIP-seq peaks (peaks detected at *P* < 1e-5 in at least one replicate) (**Table S14**). We also performed a motif enrichment analysis for the target CpGs of rs1223592 *trans*-meQTL using PWMEnrich, which showed that the motifs for IRF family proteins ranked at the top, with the IRF4 motif being the second most significantly enriched motif (*P* = 3.09e-14) (**Table S16; Fig S12**). We further examined differentially expressed genes in 501mel cells with *IRF4* knockdown. Among 804 differentially expressed genes upon IRF4 knockdown (*P* < 0.01 and |log2(fold change)| > 1), 7 genes overlapped with 8 target CpG probes of rs12203592 *trans*-meQTL (*VPS13B, NCKAP5, E2F5, RGMB, SMG6, MYH10, MAP2K6*) (Enrichment OR = 2.8, *P*-value = 0.1), while none of them are near ChIP-seq peaks. These results indicate that *IRF4* is a melanocyte-specific transcriptional regulator of multiple target genes that are under tight genetic control. The data also supports the hypothesis that allelic methylation changes in *trans* reflect altered gene expression driven by transcription factor binding, rather than methylation changes themselves driving expression changes.

Finally, we tested if significant melanocyte *trans*-meQTLs were also present in melanomas. Among 15,179 *trans*-meQTL variant-meProbe pairs found in melanocytes (*FDR* < 0.01), 11,714 were present in TCGA SKCM dataset. rs12203592 was not present in the TCGA dataset and could not be tested. Of the tested variant-meProbe pairs, 9,868 (65% of 15,179 or 84% of 11,714) including all 332 melanocyte *trans*-meProbes were significant in melanomas (*P* < 1e-11; equivalent to *FDR* < 0.01). A strong correlation of *trans*-meQTL effect sizes was observed between melanocyte and melanoma datasets (Pearson R = 0.71; *P* < 2.2e-16) (**Fig S13**). These data indicated that melanocyte *trans*-meQTLs are highly preserved in malignant melanomas.

## Discussion

To date, meQTL studies have been mainly performed in blood and blood cell types^11, 12, 14–20^, tumor tissues^24, 25^, and/or normal bulk tissues^13, 21–23^. However, cell-type-specific meQTL studies using the cell of origin for many diseases and traits have been largely lacking. Our study presents a rare example of a single cell type meQTL dataset accompanied by matching expression QTL data. In this study, we explored the roles of cell-type-specific meQTL in characterizing disease-associated genomic variants as well as understanding their roles in gene expression regulation. Using multi-trait colocalization and MWAS, we demonstrated that a melanocyte meQTL generated from a dataset of moderate sample size (n = 106) provides substantial power to detect melanoma-associated CpG probes. Comparison of meQTLs between melanocytes and malignant melanomas revealed that melanocyte meQTLs are far better preserved than eQTLs in melanomas. Together, melanocyte multi-QTL and melanoma meQTL nominated molecular phenotypes underlying 72% of known genome-wide significant melanoma GWAS loci (and identified multiple novel loci), which is higher than conventional eQTL colocalization-based findings^1^. Pathway analyses of these genes highlighted melanoma- and melanocyte lineage-specific signaling, as well as a master regulator of melanocyte lineage, MITF, which was not apparent from the analyses using only eQTL. Melanocyte meQTL also extended our knowledge on genetic regulation of gene expression involving DNA methylation. eQTL-meQTL colocalization/mediation analyses and *trans*-meQTL hotspot analysis highlighted the roles of transcription factors in allelic methylation patterns including those through lineage-specific transcription factors and target genes.

Melanocyte *trans*-meQTL analysis identified a melanocyte-specific regulatory network involving a transcription factor, IRF4. Previous studies suggested that *trans*-meQTL hotspots could affect the expression of nearby transcription factors (i.e. *cis*-eQTL), which might be reflected on the allelic methylation of their potential binding sites across the genome^11–13^. In our study, a *trans*-meQTL hotspot SNP, rs12203592, displayed multiple lines of support for regulation by the IRF4 transcription factor. IRF4 is primarily known as an interferon regulatory factor highly expressed in lymphocytes and blood cells, but rs12203592 is located in a melanocyte-specific enhancer element and seems to be regulated through a melanocyte-lineage specific transcriptional program affecting pigmentation phenotypes^54^. Consistent with this observation, two large blood *trans*-meQTL studies using thousands of samples did not identify *trans*-meQTL hotspots through rs12203592^11, 12^. Among the target CpGs of rs12203592 *trans*-meQTLs is the recently identified melanoma susceptibility gene, *MX2*, which also has pleiotropic roles in both melanoma promotion and immune response, hinting at potential functional interaction between *IRF4* and *MX2* in melanomagenesis. By combining eQTL, meQTL, and mediation analysis as well as ChIP-seq and knockdown analyses, our study presents a unique example of a cell-type-specific transcriptional network mediated by a multi-function transcription factor. Notably, the IRF4-mediated regulatory network in melanocytes was marginally detectable by *trans*-eQTL^6^, but *trans*-meQTL analysis in the current study revealed orders of magnitude larger plausible downstream targets (4 genes at *FDR* < 0.1 vs. 131 CpGs at *FDR* < 0.01). These data suggest that CpG methylation might better represent the dynamic status of transcription factor binding-related chromatin changes than gross gene expression changes do.

Our study provides one of the few formal comparisons of meQTLs and eQTLs between tumor tissues and cells of tumor origin. We show that a substantial proportion (45.9%) of genome-wide significant meCpG probes in melanocytes are preserved in melanomas. This is a much larger overlap compared to that of eGenes observed in our previous eQTL study using the same datasets, where only 12.7% of melanocyte eGenes were preserved in TCGA melanomas. Loss of the majority of normal tissue eQTLs in tumors was also observed in prostate tumors although this was not examined genome-wide^21^. Our comparisons of eQTL and meQTL from the same samples suggest that genetic control of lineage-specific CpG methylation is largely still detectable even in the presence of presumably high variation of methylation in tumor genomes. Our eQTL-meQTL colocalization analysis also indicated that a substantial portion of tested genes in melanocytes are potentially co-regulated with DNA methylation through common genetic variants. Importantly, these co-regulated genes and CpG probes are likely to remain under genetic control during malignant transformation even in the presence of somatic events. Consistent with this idea, melanocyte *trans*-meQTLs (presumably regulated through transcription factor binding) were preserved in melanomas at an even higher level (65%) than *cis*-meQTLs. These data provide an insight into our understanding of gene expression regulation in tumors, where both heritable and tumor-specific events contribute to the total transcriptome profile.

Although meQTL is powerful, sensitive, and reliable, assigning the effector genes to significant meCpG is still challenging in the absence of colocalizing eQTL support. Colocalization approaches with an improved detection power might help identify those left undetected with the current approaches. Additionally, some of the GWAS-colocalizing meQTLs without concurrent eQTL support might reflect loci poised to be connected with allelic differences in gene expression upon proper stimulations (e.g. UV exposure), which actively proliferating cultured melanocytes cannot recapitulate.

In conclusion, our study demonstrated the utility of cell-type-specific meQTL in GWAS annotation and provided insights into melanocyte-specific gene expression regulation involving DNA methylation.

## Supplemental Data

Supplemental Data include thirteen figures and sixteen tables.

## Acknowledgments

This work has been supported by the Intramural Research Program (IRP) of the Division of Cancer Epidemiology and Genetics, National Cancer Institute, US National Institutes of Health. This work was also supported by grants from the Research Fund of Iceland to ES (184861 and 207067), a grant from the University of Iceland Doctoral Grants Fund to RD, and a postdoctoral grant from the University of Iceland Research Fund to BOE, as well as funding from Cancer Research UK (programme award C588/A19167) and by the NIH (R01 CA83115). Genotyping services were provided by the Center for Inherited Disease Research (CIDR). CIDR is fully funded through a federal contract from the National Institutes of Health to The Johns Hopkins University, contract number HHSN268201100011I. This work utilized the computational resources of the NIH high-performance computational capabilities Biowulf cluster (http://hpc.nih.gov). The results appearing here are in part based on data generated by the TCGA Research Network (http://cancergenome.nih.gov/). We would like to thank members at the National Cancer Institute Cancer Genomics Research Laboratory (CGR) for help with sequencing efforts, Stacie Loftus and William Pavan from National Human Genome Research Institute for the help with the melanocyte eQTL study, and Christopher Foley from the University of Cambridge for the help with HyPrColoc analysis. We also thank all the cohorts, funders, and investigators who contributed to the melanoma GWAS, acknowledged by Landi, 23andMe, and colleagues, from which data was used towards fine-mapping. We would like to thank the research participants and employees of 23andMe for making this work possible. The content of this publication does not necessarily reflect the views or policies of the US Department of Health and Human Services, nor does mention of trade names, commercial products, organizations imply endorsement by the US government.

## Web Resources

minfi: https://bioconductor.org/packages/release/bioc/html/minfi.html

FastQTL: http://fastqtl.sourceforge.net/

LeafCutter: https://davidaknowles.github.io/leafcutter/

tensorQTL: https://github.com/broadinstitute/tensorqtl

QVALUE: https://bioconductor.org/packages/release/bioc/html/qvalue.html

HyPrColoc: https://github.com/jrs95/hyprcoloc

TWAS FUSION: http://gusevlab.org/projects/fusion/

eQTLMAPT: https://github.com/QidiPeng/eQTLMAPT

PLINK: https://www.cog-genomics.org/plink/

PWMEnrich: https://bioconductor.org/packages/PWMEnrich/

epitools: https://CRAN.R-project.org/package=epitools

MACS: https://github.com/macs3-project/MACS

Kallisto: https://pachterlab.github.io/kallisto/

Sleuth: https://pachterlab.github.io/sleuth/about

Haploreg: https://pubs.broadinstitute.org/mammals/haploreg/haploreg.php

GDC Data Portal: https://portal.gdc.cancer.gov

Ingenuity Pathway Analysis: http://www.ingenuity.com/index.html

## Data and Code Availability

The raw data of Illumina HumanMethylation450 BeadChip from 106 primary human melanocytes have been submitted to the Gene Expression Omnibus (GEO) database under accession code GSE166069; Melanocytes genotype data, RNA-Seq expression data, and all meQTL association results are deposited in Genotypes and Phenotypes (dbGaP) under accession phs001500.v1.p1. IRF4 ChIPseq and RNAseq data are deposited in GEO under accession code GSE167945. Data from the 2020 melanoma GWAS meta-analysis performed by Landi and colleagues were obtained from dbGaP (phs001868.v1.p1), with the exclusion of self-reported data from 23andMe and UK Biobank. The full GWAS summary statistics for the 23andMe discovery data set will be made available through 23andMe to qualified researchers under an agreement with 23andMe that protects the privacy of the 23andMe participants. Please visit https://research.23andme.com/collaborate/#dataset-access/ for more information and to apply to access the data. Summary data from the remaining self-reported cases are available from the corresponding authors of that manuscript (Matthew Law; Mark Iles; and Maria Teresa Landi).

