## Supplemental Figures for "Cell-type-specific meQTL extends melanoma GWAS annotation beyond eQTL and informs melanocyte gene regulatory mechanisms"

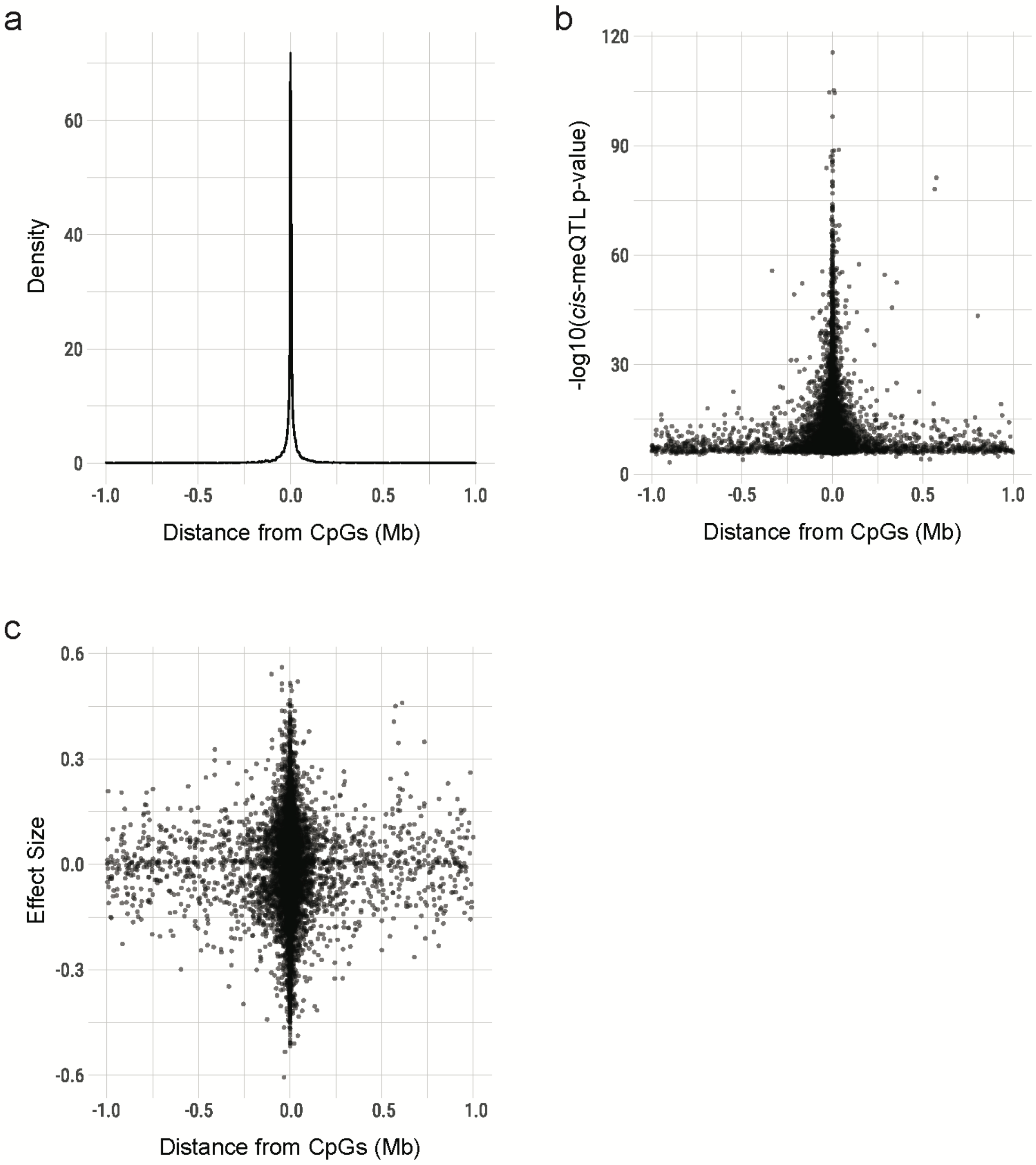
a
Density
60
40
20
0
-1.0
-0.5
0.0
0.5
1.0
Distance from CpGs (Mb)
b
-log10(cis-meQTL p-value)
120
90
60
30
0
-1.0
-0.5
0.0
0.5
1.0
Distance from CpGs (Mb)
c
Effect Size
0.6
0.3
0.0
-0.3
-0.6
-1.0
-0.5
0.0
0.5
1.0
Distance from CpGs (Mb)

**Figure S1**

**Figure S1. Melanocyte *cis*-meQTL SNPs are enriched near CpG sites.** (a) Density, (b) -$log_{10}$(meQTL P-values), and (c) meQTL effect size are plotted against distance from the CpG nearest sites in a +/- 1Mb window relative to each CpG site for each melanocyte meQTL SNP. Each dot represents significant eQTL SNP based on FDR < 5%.

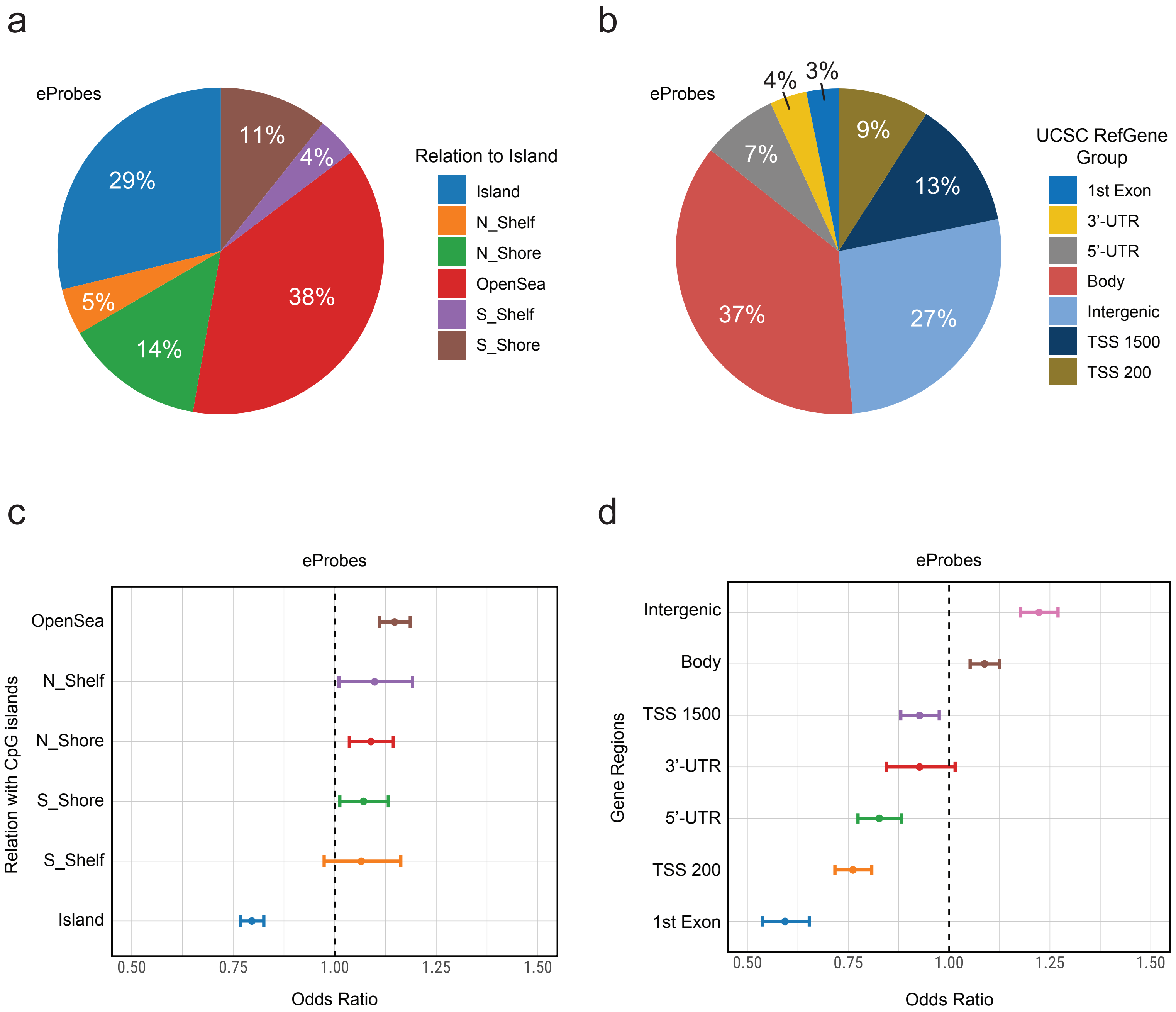
a
ePobes
29%
11%
4%
38%
14%
5%
Relation to Island
Island
N_Shelf
N_Shore
OpenSea
S_Shelf
S_Shore
b
ePobes
4%
3%
9%
7%
13%
37%
27%
UCSC RefGene Group
1st Exon
3'-UTR
5'-UTR
Body
Intergenic
TSS 1500
TSS 200
c
eProbes
OpenSea
N_Shelf
N_Shore
S_Shore
S_Shelf
Island
Relation with CpG islands
0.50
0.75
1.00
1.25
1.50
Odds Ratio
d
eProbes
Intergenic
Body
TSS 1500
3'-UTR
5'-UTR
TSS 200
1st Exon
Gene Regions
0.50
0.75
1.00
1.25
1.50
Odds Ratio

Figure S2

**Figure S2. Characteristics of melanocyte *cis*-meQTL probes.** Pie charts show the distributions of genome-wide melanocyte meQTL probes based on their genomic position in relation to CpG islands (a) and nearby genes (b)**.** Forest plots show enrichment or depletion of significant meQTL probes in each catalog in relation to CpG islands (c) and gene region (d); Odds ratios with 95% confidence level are calculated by comparing overlaps between each feature and significant meQTLs probes determined by chance alone.

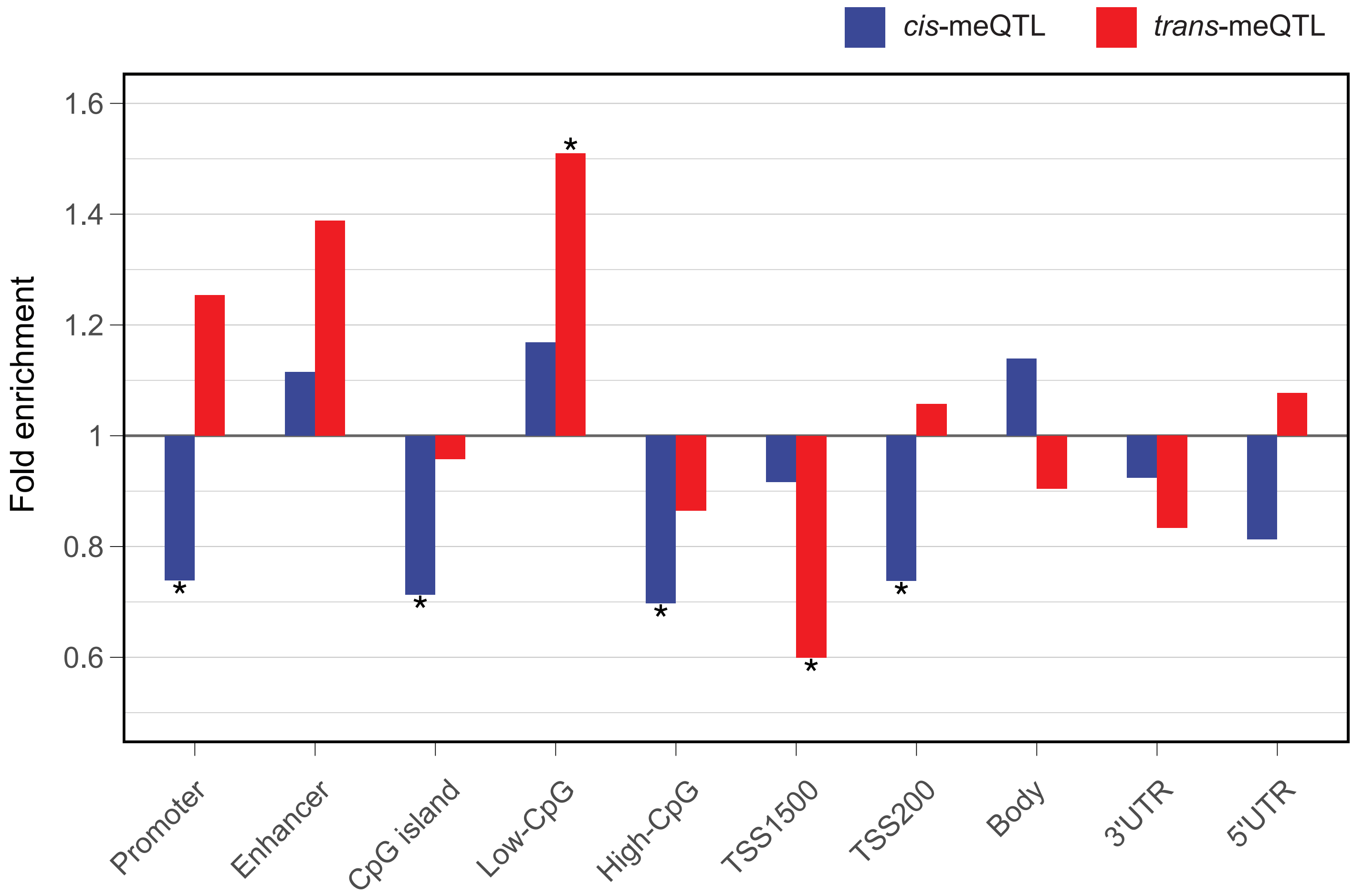

cis-meQTL
trans-meQTL
Fold enrichment
1.6
1.4
1.2
1
0.8
0.6
*
Promoter
Enhancer
CpG island
Low-CpG
High-CpG
TSS1500
TSS200
Body
3'UTR
5'UTR


Figure S3

**Figure S3. Genomic features enrichment for melanocyte meQTL probes.** Enrichment of *cis*- (blue) and *trans* (red) -meQTLs CpGs (meCpGs) in different genomic regions. Asterisk Indicates significant results with fold change > 1.2 or < 0.8, and P < 0.05/10.

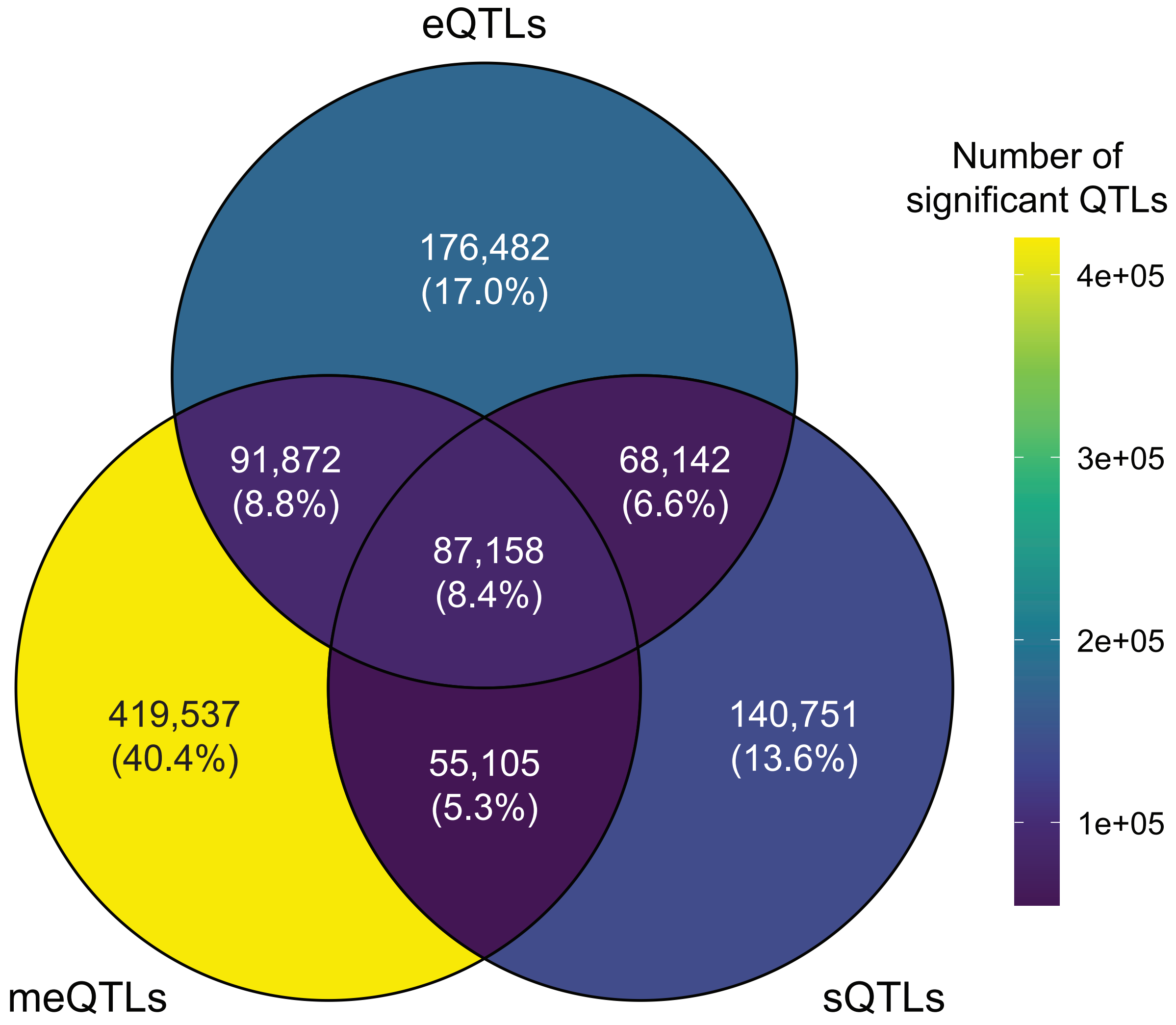
eQTLs
meQTLs
sQTLs
176,482
(17.0%)
91,872
(8.8%)
68,142
(6.6%)
87,158
(8.4%)
419,537
(40.4%)
55,105
(5.3%)
140,751
(13.6%)
Number of
significant QTLs
4e+05
3e+05
2e+05
1e+05

Figure S4

**Figure S4. Venn diagram of overlapping SNPs among melanocyte genome-wide significant eQTLs, meQTLs and sQTLs.**

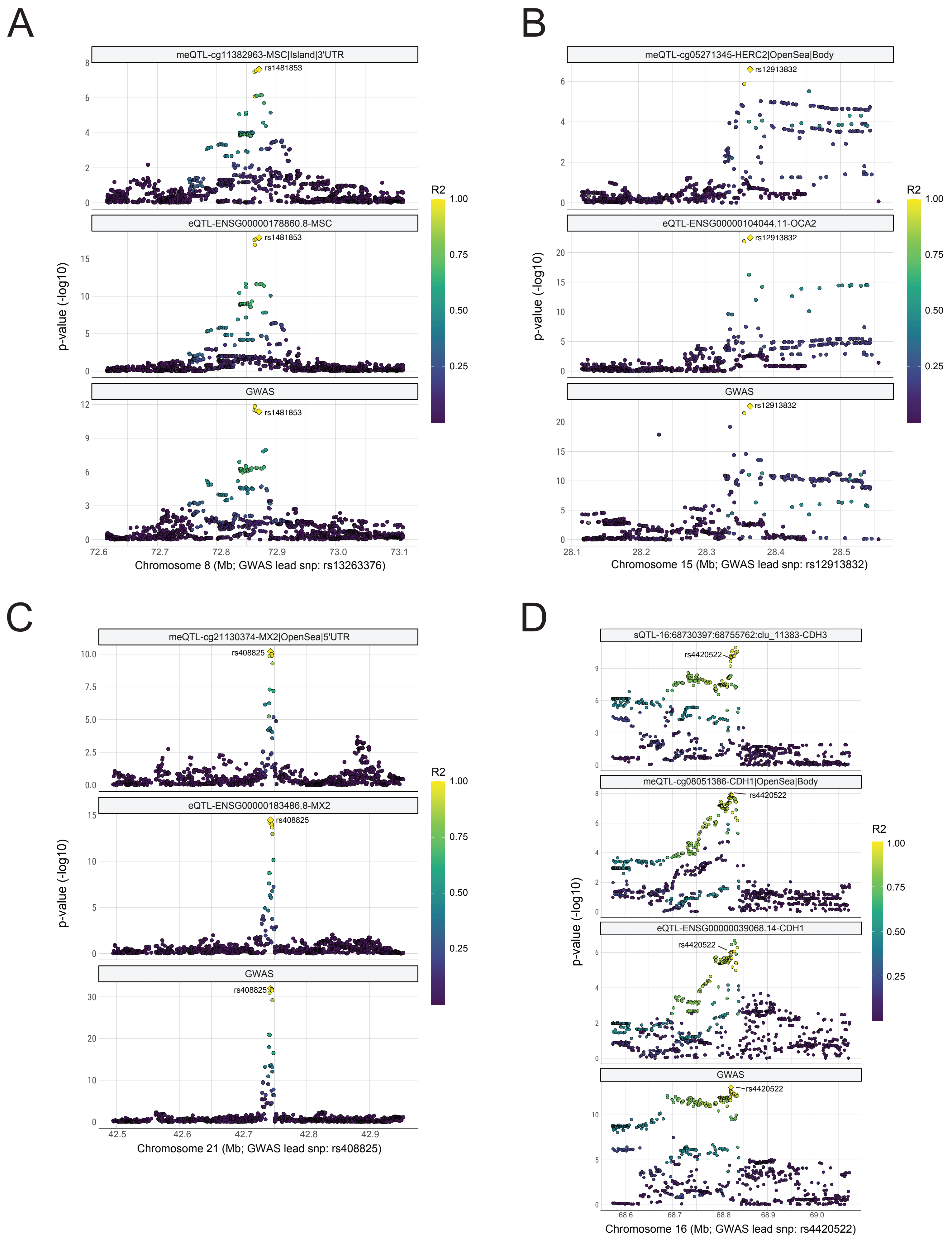
A
meQTL-cg11382963-MSC|Island|3'UTR
eQTL-ENSG00000178860.8-MSC
GWAS
rs1481853
p-value (-log10)
Chromosome 8 (Mb; GWAS lead snp: rs13263376)
B
meQTL-cg05271345-HERC2|OpenSea|Body
eQTL-ENSG00000104044.11-OCA2
GWAS
rs12913832
p-value (-log10)
Chromosome 15 (Mb; GWAS lead snp: rs12913832)
C
meQTL-cg21130374-MX2|OpenSea|5'UTR
eQTL-ENSG00000183486.8-MX2
GWAS
rs408825
p-value (-log10)
Chromosome 21 (Mb; GWAS lead snp: rs408825)
D
sQTL-16:68730397:68755762:clu_11383-CDH3
meQTL-cg08051386-CDH1|OpenSea|Body
eQTL-ENSG00000039068.14-CDH1
GWAS
rs4420522
p-value (-log10)
Chromosome 16 (Mb; GWAS lead snp: rs4420522)
R2
1.00
0.75
0.50
0.25

Figure S5

**Figure S5. Multi-QTL colocalizations were identified in four loci where more than one cell-type specific QTL trait colocalizes with the melanoma GWAS signal.** Locuszoom plots show the -log10(p-value) from GWAS meta-analysis and QTL association. GWAS lead SNPs are labeled in each locuszoom plot and the LD to GWAS lead SNPs are calculated based on 1000 genome EUR dataset. (A) *MSC/RP11-383H13.1,* (B) *OCA2/AC090696.2,* (C) *MX2*, (D) *CDH1*/*CDH3.* (A-C) SNP-Gene-meCpG trios with the highest posterior probability for each locus are shown.

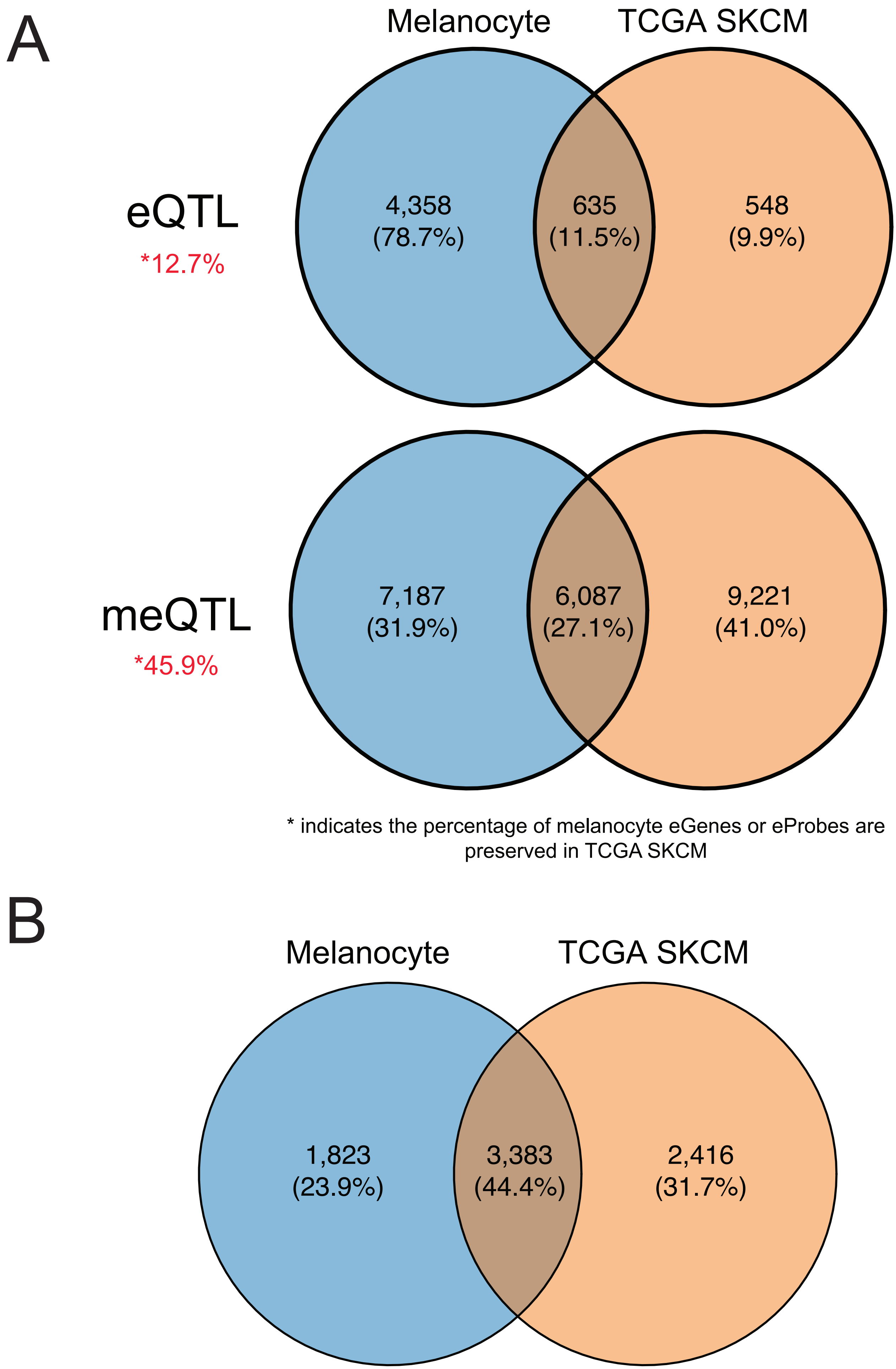
A
Melanocyte
TCGA SKCM
eQTL
*12.7%
4,358
(78.7%)
635
(11.5%)
548
(9.9%)
meQTL
*45.9%
7,187
(31.9%)
6,087
(27.1%)
9,221
(41.0%)
* indicates the percentage of melanocyte eGenes or eProbes are preserved in TCGA SKCM
B
Melanocyte
TCGA SKCM
1,823
(23.9%)
3,383
(44.4%)
2,416
(31.7%)

Figure S6

**Figure S6. Melanocyte meQTLs are preserved in melanomas.** (A) Comparisons of normal to tumor preservation between melanocyte eGenes and meProbes. (B) Preservation of melanocyte meQTL in melanomas at the gene levels. When meProbes were assigned to nearby genes based on Illumina annotation, 65% of melanocyte meQTLs were preserved in melanomas at the gene level.

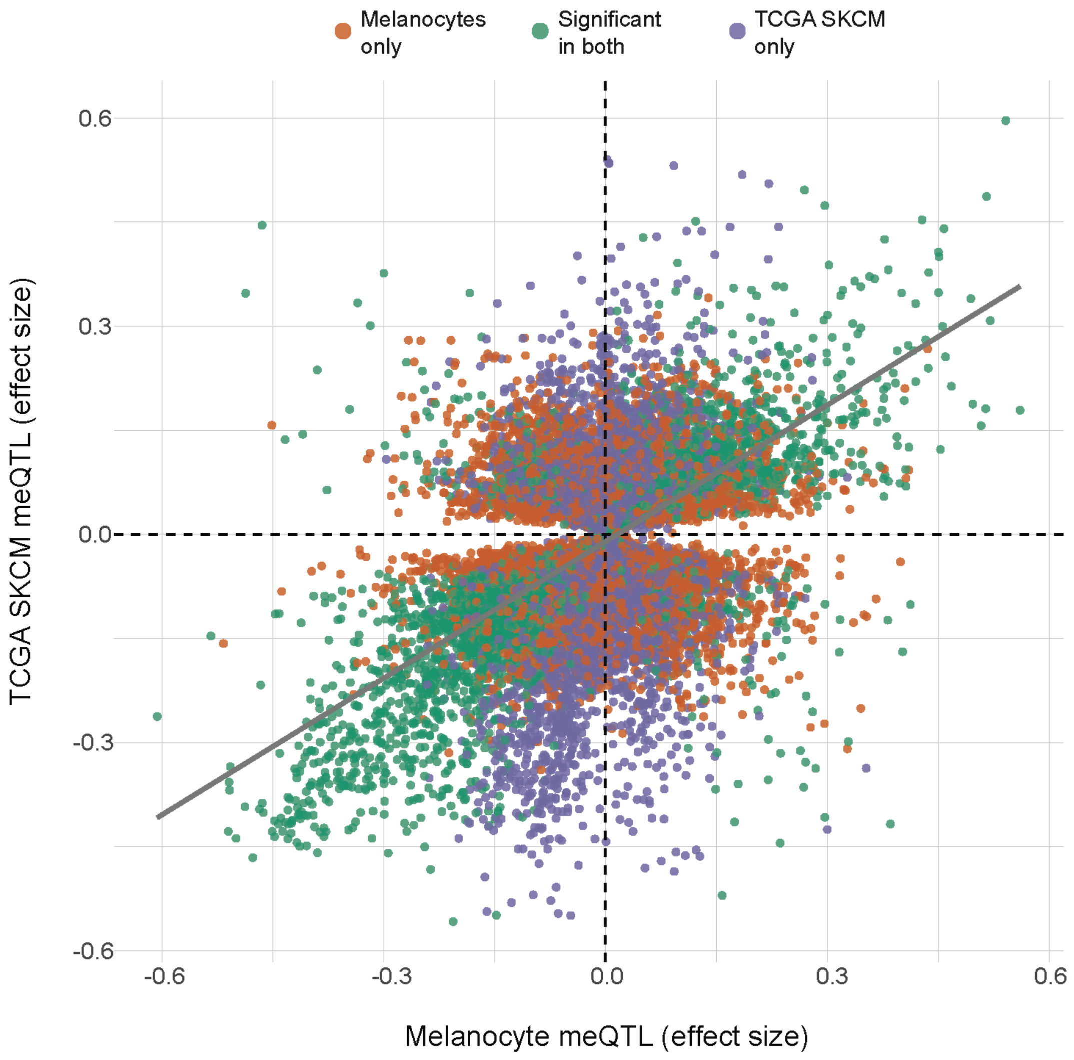

Melanocytes only
Significant in both
TCGA SKCM only
TCGA SKCM meQTL (effect size)
0.6
0.3
0.0
-0.3
-0.6
-0.6
-0.3
0.0
0.3
0.6
Melanocyte meQTL (effect size)


**Figure S7**

**Figure S7. Comparison of meQTL effect size between melanocyte and melanoma datasets.** The colors represent the overlapping of genome wide significance of meQTLs in melanocyte and melanoma datasets.

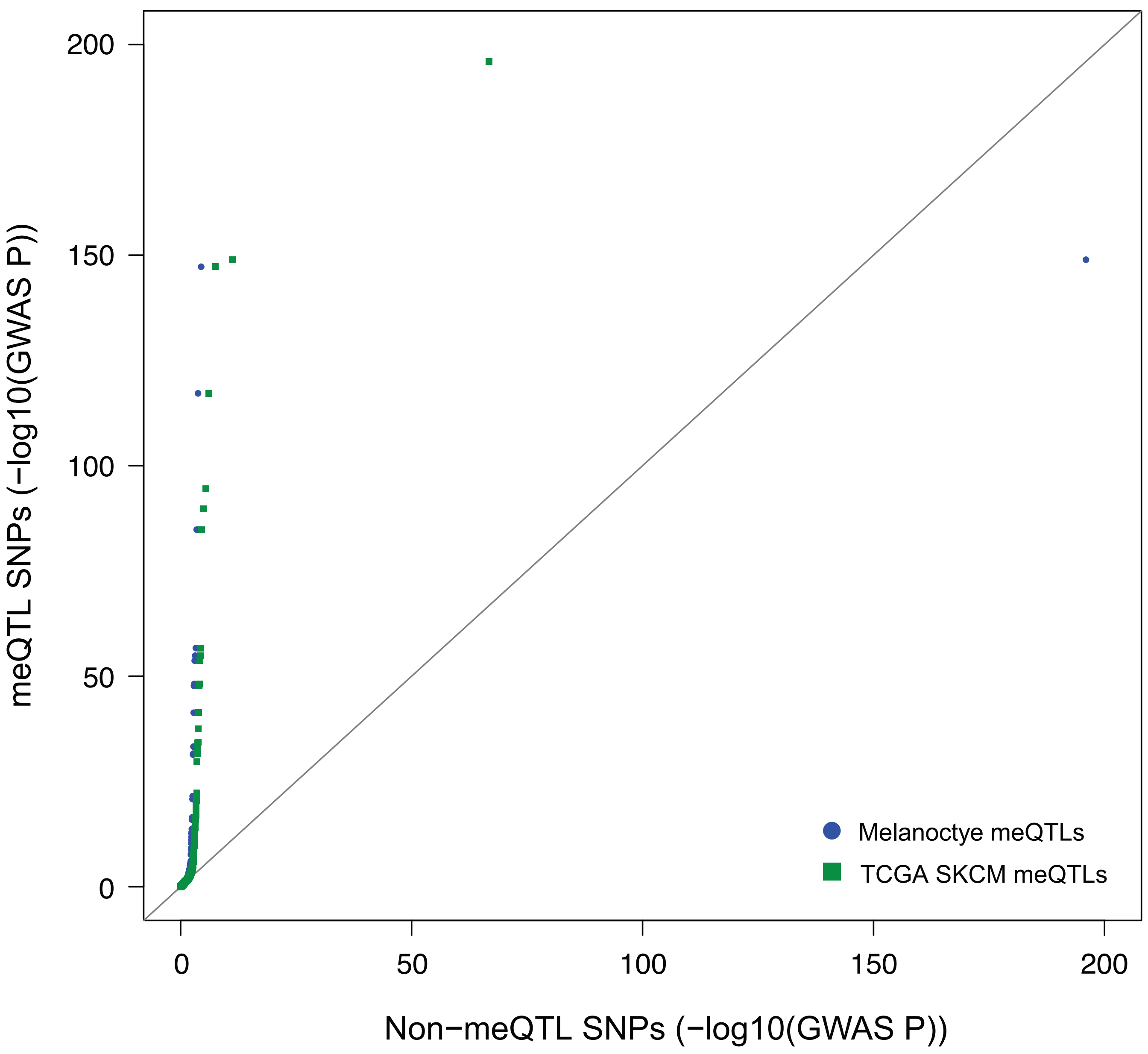

meQTL SNPs (−log10(GWAS P))
200
150
100
50
0
Melanoctye meQTLs
TCGA SKCM meQTLs
0
50
100
150
200
Non−meQTL SNPs (−log10(GWAS P))


Figure S8

**Figure S8. Enrichment of melanocytes and melanomas *cis*-meQTL variants in melanoma risk-associated variants.** QQ plot presents melanoma GWAS LD-pruned *p*-values of significant meQTL SNPs versus non-meQTL SNPs for the melanocyte data set compared to those for melanoma tumors. SNPs were classified as meQTL SNPs if there were significant meQTLs or in strong LD ($r^2$>0.8) with an meQTL SNP in each data set.

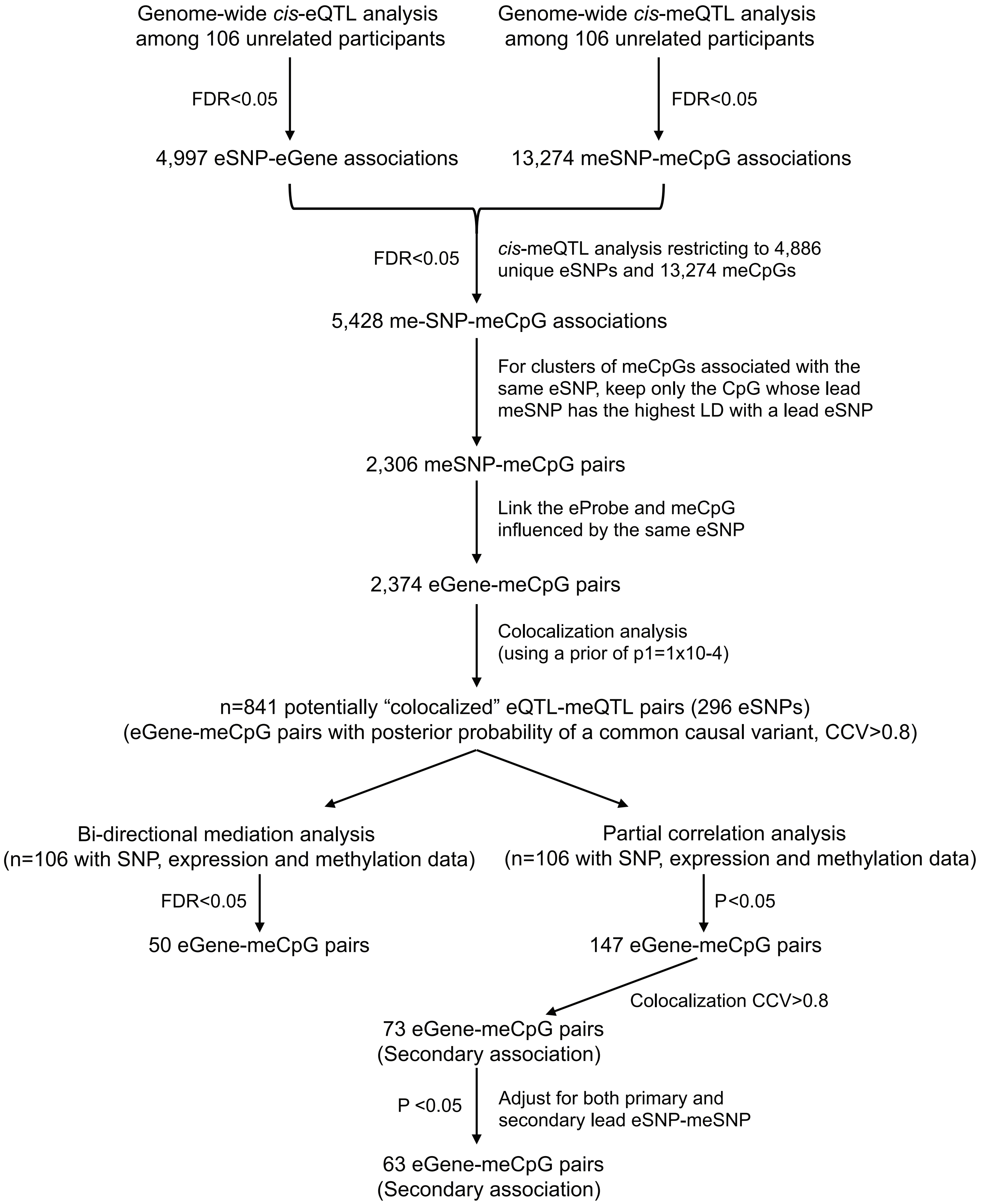
Genome-wide *cis*-eQTL analysis among 106 unrelated participants
Genome-wide *cis*-meQTL analysis among 106 unrelated participants
FDR<0.05
FDR<0.05
4,997 eSNP-eGene associations
13,274 meSNP-meCpG associations
FDR<0.05
*cis*-meQTL analysis restricting to 4,886 unique eSNPs and 13,274 meCpGs
5,428 me-SNP-meCpG associations
For clusters of meCpGs associated with the same eSNP, keep only the CpG whose lead meSNP has the highest LD with a lead eSNP
2,306 meSNP-meCpG pairs
Link the eProbe and meCpG influenced by the same eSNP
2,374 eGene-meCpG pairs
Colocalization analysis (using a prior of p1=1x10-4)
n=841 potentially “colocalized” eQTL-meQTL pairs (296 eSNPs)
(eGene-meCpG pairs with posterior probability of a common causal variant, CCV>0.8)
Bi-directional mediation analysis (n=106 with SNP, expression and methylation data)
Partial correlation analysis (n=106 with SNP, expression and methylation data)
FDR<0.05
P<0.05
50 eGene-meCpG pairs
147 eGene-meCpG pairs
Colocalization CCV>0.8
73 eGene-meCpG pairs (Secondary association)
P <0.05
Adjust for both primary and secondary lead eSNP-meSNP
63 eGene-meCpG pairs (Secondary association)

Figure S9

**Figure S9. Workflow of mediation and partial correlation analysis using both *cis*-meQTL and *cis*-eQTL dataset**

a

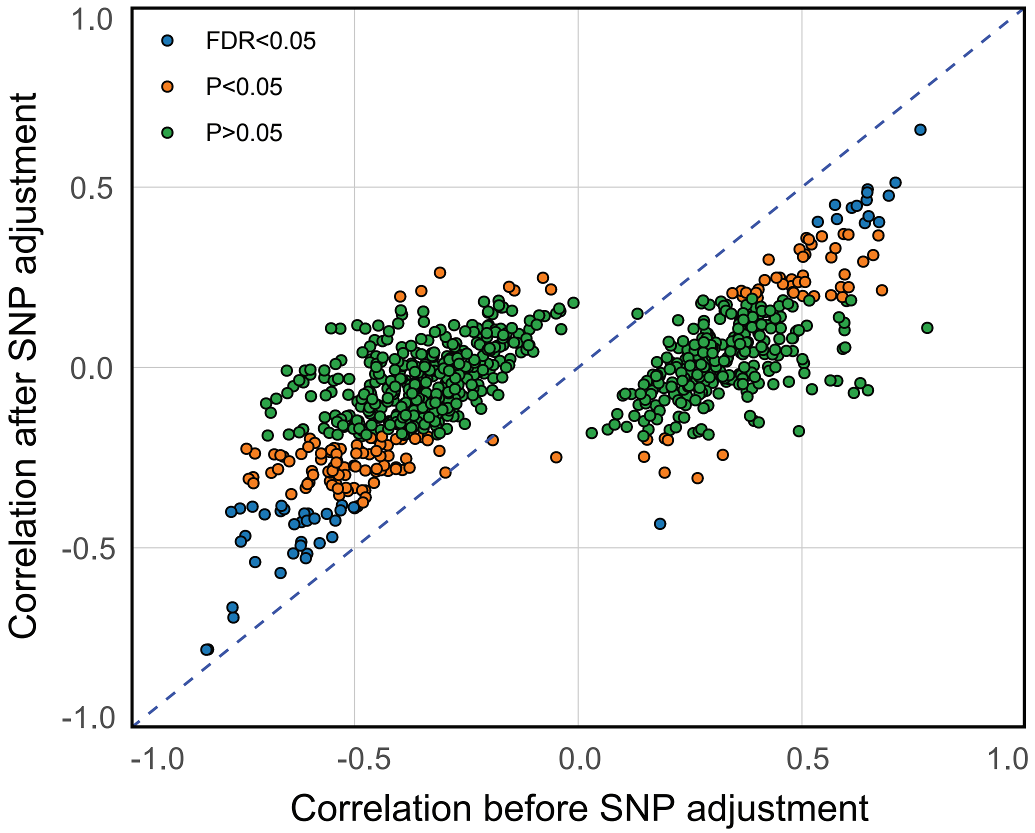


b

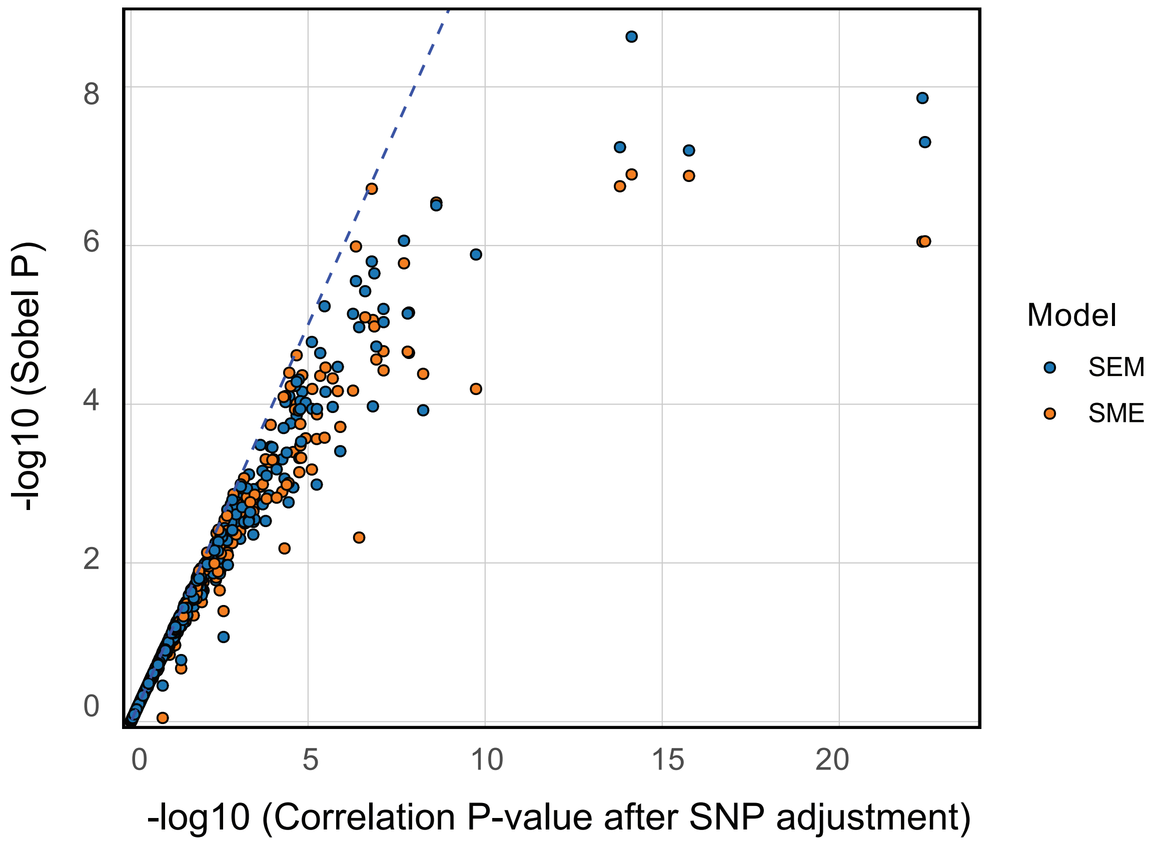


Figure S10

**Figure S10. Partial correlation and mediation analysis of 841 potentially colocalizing SNP-meProbe-eGene trios in melanocytes.** (A) correlation between eGene and meProbe before and after adjusting for the SNP, (B) relationship between the post-adjustment correlation P values from partial correlation analysis and Sobel P from mediation analyses.

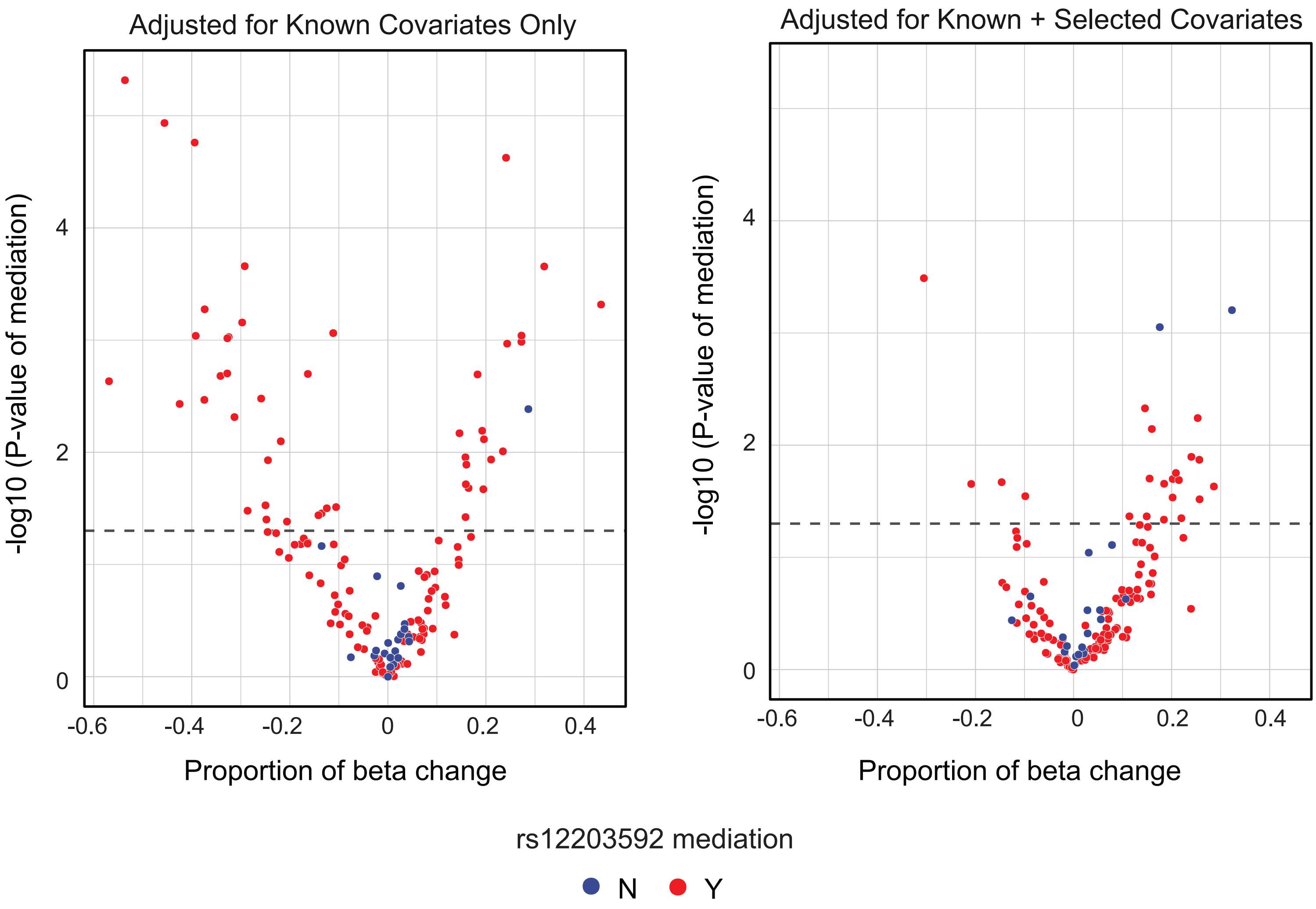
Adjusted for Known Covariates Only
Adjusted for Known + Selected Covariates
-log10 (P-value of mediation)
Proportion of beta change
-0.6
-0.4
-0.2
0
0.2
0.4
0
2
4
rs12203592 mediation
N
Y

Figure S11

**Figure S11.** Mediation analysis for melanocyte *cis*-eQTL and *trans*-meQTL. The p-values are calculated based on (left) mediation tests without adjusting for hidden confounders (right) mediation tests by eQTLMAPT considering all methylation PEER factors as potential confounders. Trios involving the *trans*-meQTL rs12203592 are highlighted in red.

| Rank | Target | PWM | Motif ID | Raw | P-value | Rank |
|---|---|---|---|---|---|---|
| 1 | IRF8 | 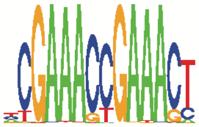 | Hsapiens−jolma2013−IRF8−2 | 15.2 | 6.83e−16 | 16 % |
| 2 | IRF4 | 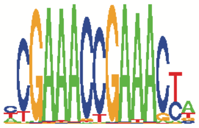 | Hsapiens−jolma2013−IRF4 | 8.65 | 3.09e−14 | 15 % |
| 3 | IRF5 | 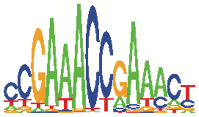 | Hsapiens−jolma2013−IRF5 | 18.1 | 4.03e−13 | 13 % |
| 4 | IRF5 | 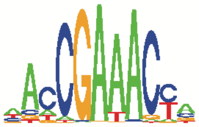 | Hsapiens−jolma2013−IRF5−2 | 5.35 | 9.9e−13 | 19 % |
| 5 | IRF8 | 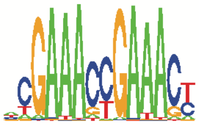 | Hsapiens−jolma2013−IRF8 | 39.2 | 2.62e−11 | 15 % |
| 6 | IRF9 | 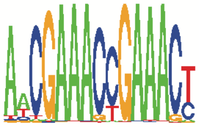 | Hsapiens−jolma2013−IRF9 | 28.6 | 3.09e−10 | 13 % |
| 7 | IRF2 | 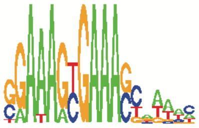 | IRF2 | 41.3 | 8.8e−10 | 15 % |
| 8 | IRF1 | 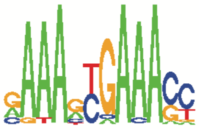 | IRF1 | 28.3 | 1.23e−09 | 18 % |
| 9 | SOX14 | 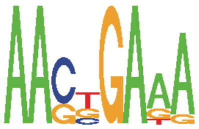 | SOX14 | 5.37 | 2.78e−09 | 15 % |
| 10 | STAT2::STAT1 | 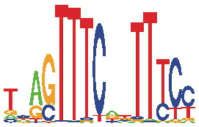 | STAT2::STAT1 | 219 | 7.09e−09 | 21 % |

**Figure S12**

**Figure S12.** Motifs enriched near the target CpGs of IRF4 *trans*-meQTLs rs12203592. The top 10 enriched motifs from PWMEnrich analysis include the rank, target, PWM sequence logo, motif id, raw score, *p*-value and the frequency.

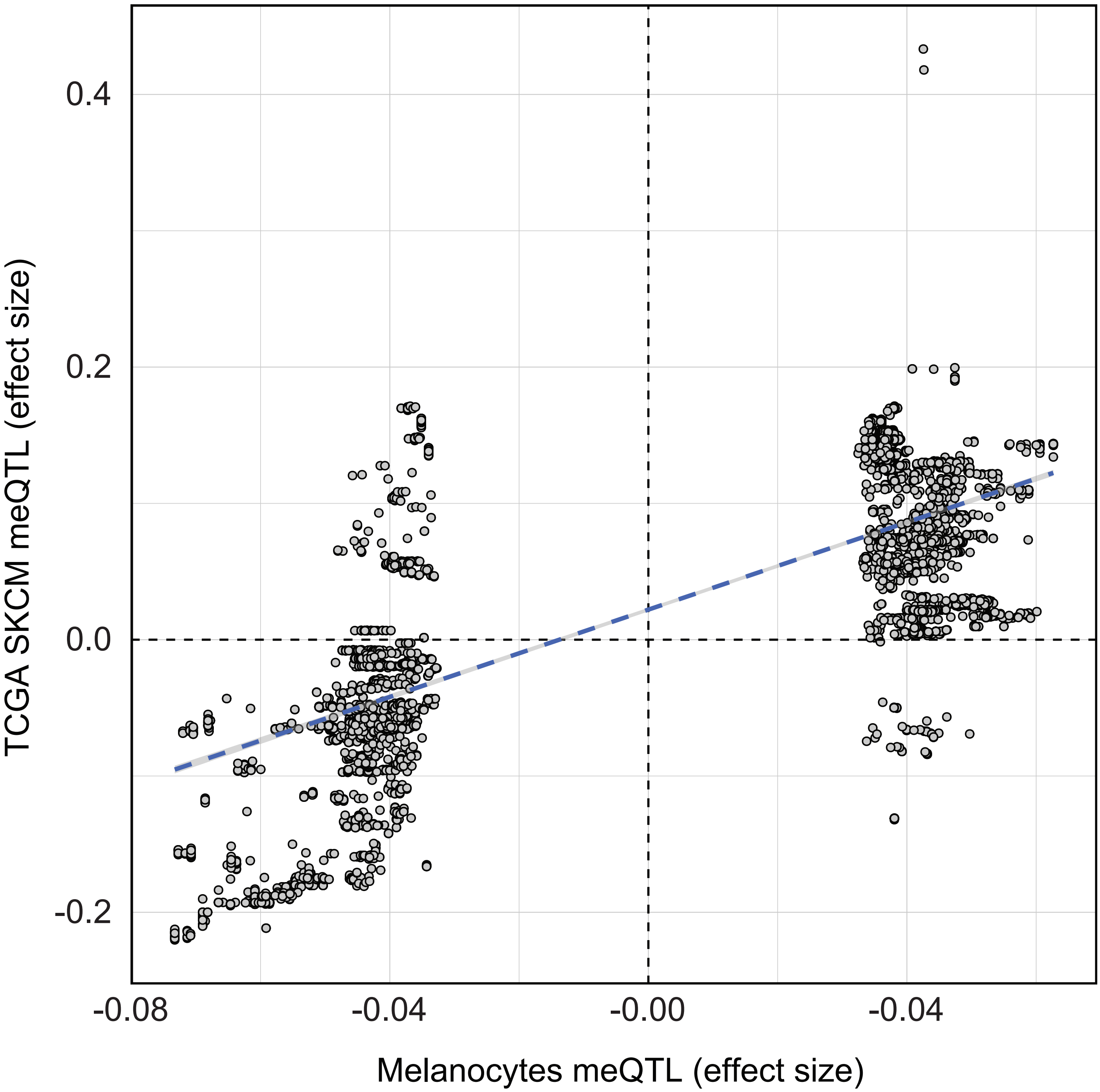
0.4
0.2
0.0
-0.2
TCGA SKCM meQTL (effect size)
-0.08
-0.04
-0.00
-0.04
Melanocytes meQTL (effect size)

Figure S13

**Figure S13. Correlation of effect sizes between *trans*-meQTLs of melanocytes and TCGA melanomas.** Only genome-wide significant *trans*-meQTLs with association *P*-value < 1.03e-11 in melanocyte data are shown here.
